# View-channel-depth light-field microscopy: real-time volumetric reconstruction of biological dynamics by deep learning

**DOI:** 10.1101/432807

**Authors:** Zhaoqiang Wang, Lanxin Zhu, Hao Zhang, Guo Li, Chengqiang Yi, Yi Li, Yicong Yang, Yichen Ding, Mei Zhen, Shangbang Gao, Tzung K. Hsiai, Peng Fei

**Affiliations:** School of Optical and Electronic Information-Wuhan National Laboratory for Optoelectronics, Huazhong University of Science and Technology, Wuhan, 430074, China; College of Life Science and Technology, Huazhong University of Science and Technology, Wuhan, 430074, China; Department of Bioengineering, Henry Samueli School of Engineering and Applied Science, University of California, Los Angeles, Los Angeles, 90095, U.S.A; Division of Cardiology, Department of Medicine, David Geffen School of Medicine, University of California, Los Angeles, Los Angeles, 90095, U.S.A; Lunenfeld-Tanenbaum Research Institute, Mount Sinai Hospital, University of Toronto, Toronto, M5G 1×5, Canada; Division of Cardiology, Department of Medicine, Greater Los Angeles VA Healthcare System, Los Angeles, U.S.A

## Abstract

Light-field microscopy has emerged as a technique of choice for high-speed volumetric imaging of fast biological processes. However, artefacts, non-uniform resolution, and a slow reconstruction speed have limited its full capabilities for *in toto* extraction of the dynamic spatiotemporal patterns in samples. Here, we combined a view-channel-depth (VCD) neural network with light-field microscopy to mitigate these limitations, yielding artefact-free three-dimensional image sequences with uniform spatial resolution and three-order-higher video-rate reconstruction throughput. We imaged neuronal activities across moving *C. elegans* and blood flow in a beating zebrafish heart at single-cell resolution with volume rates up to 200 Hz.

## Introduction

A recurring challenge in biology is the quest to extract ever more spatiotemporal information from targets, as many millisecond transient cellular processes occur in three-dimensional (3D) tissues and across long time scales. Several imaging techniques, including epifluorescence and plane illumination methods, can image live samples in three dimensions at high spatial resolution^1–5^. However, they require recording a number of two-dimensional (2D) images to create a 3D volume, and the temporal resolution is compromised by the extended acquisition time of the camera.

Recently, light-field microscopy (LFM) has become the technique of choice for instantaneous volumetric imaging^6–13^. It permits the acquisition of transient 3D signals via post-processing of the light-field information recorded by a single 2D camera snapshot. Because LFM provides high-speed volumetric imaging limited only by the camera frame rate, it has delivered promising results in various applications, such as the recording of neuronal activities^7–10^ and visualization of cardiac dynamics in model organisms^11^. Despite these advances, the generally low and non-uniform spatial resolution and the presence of reconstruction artefacts prevents its more widespread application for capturing millisecond time-scale biological processes at single-cell resolution^7, 11^. While these problems can be mitigated by optimizing the way the light field is recorded^9,12^, the extra system complexity could impede the wide dissemination of the LFM technique. Furthermore, current LFMs rely heavily on a computationally-demanding iterative recovery process that intrinsically limits the overall throughput of LFM reconstruction, compromising its potential for long time-scale applications.

Here, we propose a novel LFM strategy based on a view-channel-depth neural network processing of the light-field data, termed VCD-LFM. By light-field projection using a wave optics model^13^, we generated synthetic light-field images from high-resolution 3D images experimentally obtained beforehand, and readily paired them as input and ground-truth data for network training. The VCD network (VCD-Net) was designed to extract multiple views from these 2D light-fields, and transform them back to 3D depth images, which would be compared with the high-resolution ground truth images to guide optimization of the network. Through iteratively minimizing the loss of spatial resolution by incorporating abundant structural information from training data, this deep-learning VCD-Net could be gradually strengthened until it is capable of deducing 3D high-fidelity signals at uniform resolution across the imaging depth. In addition to a higher resolution and minimizing artefacts, once the VCD-Net is properly trained, it can deduce an image stack from a light-field measurement at a millisecond time scale, showing ultra-high throughput suited for time-lapsed video processing. We demonstrated the ability of the VCD-LFM method via imaging the motor neuron activities of L4-stage *C. elegan*s rapidly moving inside a 300 × 300 × 50 μm microfluidic chamber, at an acquisition rate of 100 Hz and processing-throughput of 13 volumes per second. This allowed us to extract the 4D spatiotemporal patterns of neuronal calcium signaling and track correlated worm behaviors at single-cell resolution, which was notably better than classic light-field deconvolution microscopy (LFDM). Furthermore, we performed *in toto* imaging of the blood flow in the beating heart of zebrafish larvae, enabling velocity tracking of blood cells and ejection fraction analysis of the heartbeat across a 250 × 250 × 150 μm volume at 200 Hz.

## Results

### Principle and performance of VCD-LFM

Our VCD-LFM involves training of a VCD convolutional neural network and its inferences based on the images obtained from LFM (**Fig. 1a, Methods, Supplementary Fig. 1, 2**). To create the data for network training, we first obtained a number of high-resolution (HR) 3D images of stationary samples using synthetic or experimental methods (**Fig. 1a, Methods**). Referring to the wave optics model of LFM^13^, we projected these HR 3D images into 2D light-field images, which were then used as the input for network training (**Fig. 1b. step 1, Supplementary Note 1**). Each synthetic light-field image was first re-arranged into different views, from which features were extracted and incorporated into multiple channels in each convolutional layer. The final output channels were then assigned to a number of planes representing different depths to generate an image stack. Using cascaded convolution layers (U-Net architecture, **Supplementary Fig. 3 and Supplementary Table 1**) to repetitively extract features, this VCD procedure generated intermediate 3D reconstructions (**Fig. 1b. step 2**). The pixel-wise mean-square-error (MSE) was counted as the loss function to indicate how different these outputs were to the ground truth images. By iteratively minimizing the loss function (**Fig. 1b. step 3**), the network was gradually optimized until it could transform the synthetic light-fields into 3D images that were sufficiently similar to the ground-truth images **(Supplementary Fig. 4, Note 2**). Following training on gigavoxels of prior data, the network was capable of instantly implementing VCD transformation of sequential light-field measurements that recorded dynamic processes, and inferring a sequence of 3D volumes at a rate up to 13 volumes s^−1^ (**Fig. 1b. step 4**).

**Figure 1.**
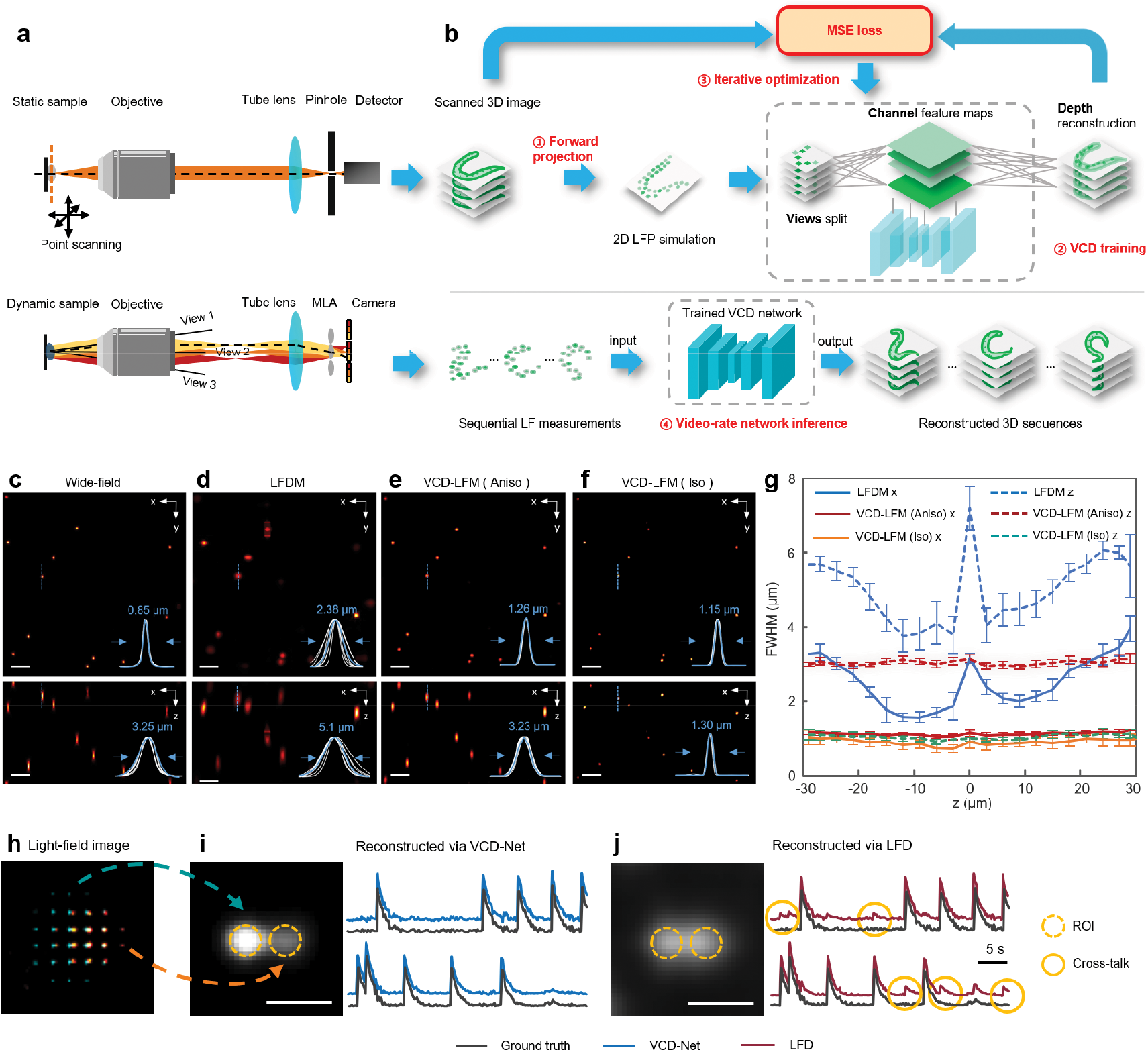
VCD-LFM and its performance. (**a**) HR 3D imaging of steady samples by a confocal microscope (upper row) and instantaneous recording of dynamic samples by a light-field microscope (bottom row). (**b**) The VCD-Net reconstruction pipeline, containing: 1. Forward light-field projection (LFP) from the HR image stacks; 2. VCD transformation of synthetic light-field inputs into intermediate 3D image stacks; 3. Network training via iteratively minimizing the difference between VCD inferences and confocal ground truths; 4. Inference of 3D images from the recorded light-field images by a well-trained VCD-Net. (**c**)**–**(**f**) Maximum intensity projections (MIPs) of the same fluorescent beads and their resolution (FWHM) by wide-field microscopy, LFDM, and VCD-LFM trained with anisotropic and isotropic HR data, respectively. White lines, intensity profiles of all the resolved beads shown in the MIPs. Blue lines, intensity profiles across a selected bead at 20 μm off the focal plane. Scale bars, 10 μm. (**g**) Average axial (dashed lines) and lateral (solid lines) FWHM of the beads across the volumes reconstructed by LFDM (n = 2,039 beads), anisotropic (n = 2,527 beads) and isotropic VCD-LFM (n = 2,731 beads), respectively. VCD-LFM improves lateral/axial resolution across the entire 60-μm overlapping volume, with Aniso-VCD achieving uniform resolution of ~1.1 ± 0.08 μm and ~3.0 ± 0.1 μm in x/y and z, respectively, and Iso-VCD achieving near isotropic resolution of ~1.0 ± 0.15 μm. Center lines represent means and error bars denote standard deviations. (**h**) One frame of a synthetic light-field video recording the activities of two adjacent firing neurons (indicated by blue and red colors). (**i**),(**j**) Reconstructions of the light-field video by VCD-Net and LFD, with signal traces extracted in the given ROIs (line circles), and compared to the ground truth. Dotted circles denote the signal cross-talks between adjacent neurons due to the blurring. Scale bar, 5 μm.

We built a custom microscope to enable *in situ* light-field recording and 3D wide-field imaging (**Methods, Supplementary Fig. 1**). To demonstrate the capability of the VCD-LFM, we reconstructed sub-diffraction beads captured using a 40×/0.8w objective, and quantified the resolution improvement resulting from the network by comparing the results with those from conventional light-field deconvolution (LFD, **Fig. 1c–f**). As verified by 3D wide-field imaging of the same volume, the fluorescence of individual beads was correctly localized throughout the volume (**Fig. 1e, f, Supplementary Fig. 5**). The VCD-LFM with a network using wide-field 3D image as the HR reference yielded averaged x,y~1.1 and z~3.0 μm resolutions (n = 2527 beads), which were also uniform across a 60-μm imaging depth (x,y~1.0 and z~2.9 μm at the best plane, x,y~1.3 and z~3.1 μm at the outer edge of the axial field of view) (**Fig. 1g, Supplementary Fig. 6**). This demonstrates significant improvements compared with x,y~2.6 and z~5.0 μm for LFDM, which are significantly more diverse (x,y~1.6 and z~3.8 μm at the best plane, x,y~4.0 and z~7.0 μm at the outer edge of the axial field of view). We note that the performance of the VCD-LFM is also training data-oriented, hence the beads can be further reconstructed isotropically (~1.0 μm) through including higher-resolution data in the training (**Fig. 1f, g, Supplementary Fig. 7).** Furthermore, VCD-LFM substantially removed the mosaic-like artefacts near the focal plane that are common in LFDM (**Fig. 1d, Supplementary Fig. 5, 6**), and it performed well even when the signals were weak with high background noise or dense with points adjacent to each other (**Supplementary Fig. 8, 9**). To further validate the accuracy of reconstructed signals, we applied VCD-Net to the reconstruction of two adjacent firing neurons. As shown in **Fig. 1h-j**, the improved image quality by VCD-Net substantively suppressed signal cross-talk by blurring and artefacts, and thus contributed to accurate recovery of signal fluctuations when recording the activities of densely labeled neurons. We further validated the high reconstruction accuracy of VCD-Net on both static and moving neurons with varying signal magnitude and density (**Supplementary Fig. 10, 11**).

### VCD-LFM revealing locomotion-associated neural activities in moving *C.elegans*

We show VCD-LFM to be suitable for capturing dynamic processes in live animals by demonstrating the imaging of neuronal activities of moving *C. elegans* and the beating heart of a zebrafish larva. A microfluidic chip was used to permit *C. elegans* (L4-stage) to rapidly move inside a micro-chamber (300 × 300× 50 μm, **Fig. 2a**). We used a 40×/0.8w objective to image the activities of motor neuron nuclei labeled with GCaMP/RFP at a 100-Hz acquisition rate, yielding 6000 light-fields in each channel in a 1-minute observation (**Method**, **Supplementary Fig. 1**). The VCD-Net provided high-quality reconstructions to visualize the neuron signaling at single-cell resolution during fast body movement (**Fig. 2b, Supplementary Fig. 12, Video 1, 2**). In contrast, LFD suffered from ambiguous cellular resolution and deteriorated image quality around the focal plane, which is a known limitation (**Fig. 2c**). We identified the specific A- and B-motor neurons that have been associated with motor-program selection, and mapped their calcium activities over time after ratiometrically correcting the GCaMP signals using RFP baselines to remove motion noises^14^ (**Fig. 2d, e, Methods, Supplementary Fig. 13, Video 3**). Meanwhile, by applying an automatic segmentation of the worm body contours (**Methods, Supplementary Note 5**), we calculated the worm’s velocity and curvatures related to its locomotion and behavior, thereby allowing classification of the worm motion into forward, backward, or irregular crawling (**Fig. 2f–h**). The patterns of transient *Ca*^2+^ signaling were found to be relevant to the switches from forward to backward crawling of the worm, which is consistent with previous findings^14–17^. Furthermore, the non-iterative VCD reconstruction could sequentially recover 3D images at a volume rate of 13.5 Hz, ~900-times faster than the iterative LFD method (**Fig. 2i, Supplementary Table 3**). Our VCD-LFM thus demonstrated significant advantages for visualizing sustained biological dynamics, which is computationally challenging using conventional deconvolution methods.

**Figure 2.**
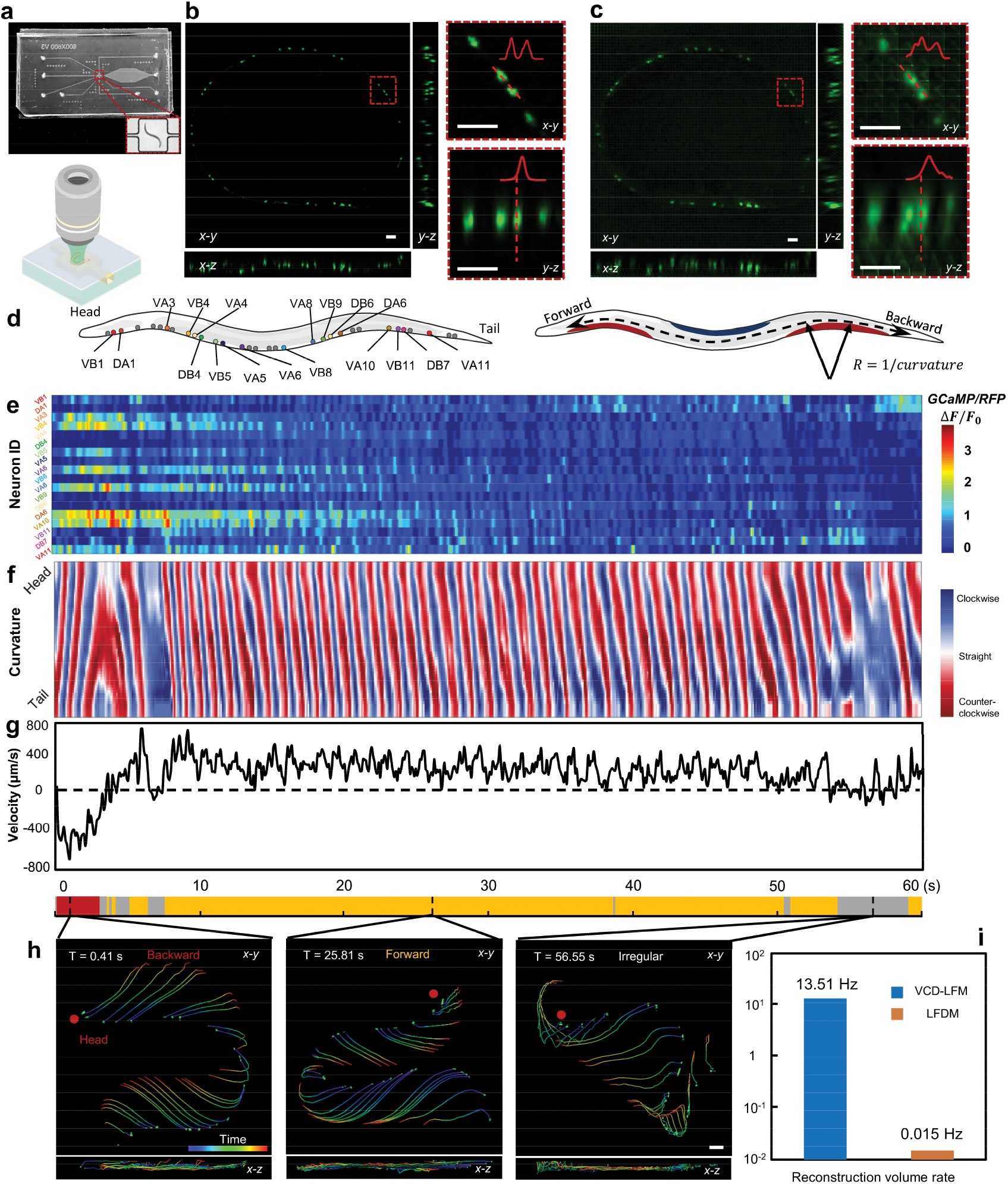
Whole-animal *Ca*^2+^ imaging of moving *C. elegans* using VCD-LFM. (**a**) Configuration combining light field with microfluidic technique for imaging motor neuron activities of L4-stage *C. elegans* (strain ZM9128 *hpIs595*[*Pacr-2(s)*::GCaMP6(f)::wCherry]) acting in a microfluidic chip (300 × 300 × 50 μm, top panel) at 100-Hz recording rate. (**b**),(**c**) MIPs of one instantaneous volume reconstructed by VCD and LFD, respectively. The magnified views of a selected region show that VCD restores single-cell resolution and removes artefacts. The data shown are representative of *n* = 10 independent *C.elegans*. Scale bars, 10 μm. (**d**) Schematics of the worm with identified motor-neurons labeled (left) and motion definition annotated (right). (**e**) Heatmap visualizing the neural activities of 18 identified motor neurons during a 1-minute observation of the acting worm. Each row shows the *Ca*^2+^ signal fluctuation of an individual neuron with color indicating the percent fluorescence changes (*ΔF/F*_*O*_), where *F* is ratiometrically corrected by the ratio of GCaMP fluorescence to RFP fluorescence. (**f**) Curvature kymograms along the body of the moving worm. (**g**) Velocity plot shows the displacement in the direction of body. An ethogram describing the worm behavior over time (lower panel) is obtained by analyzing the curvature and velocity change. (**h**) Selected volumes with time-coded traces (duration: left and middle, 150 ms; right, 500 ms) in accordance with the ethogram visualizing the backward (left), forward (middle), and irregular (right) crawling tendency of the worm. Scale bars, 20 μm. (**i**) The reconstruction throughput of VCD-LFM and LFDM for processing the same *C.elegans* light-field video.

### Volumetric imaging of fast dynamics in beating zebrafish heart

We also captured the cardiac hemodynamics in beating zebrafish heart. To reduce the background from body tissue, a laser-rod was generated to selectively illuminate the heart region^18^, and the light-field video was recorded at a 200-Hz volume rate using a 20×/0.5w objective (**Fig. 3a, Methods, Supplementary Fig. 2**). The flowing red blood cells (RBCs, *Tg(gata1a:dsRed)*) and the beating cardiomyocytes nuclei (*Tg(myl7:nls-gfp)*) were both four-dimensionally reconstructed by the VCD-Net with resolution, structure similarity, and processing throughput notably better than conventional LFD approaches (**Fig. 3b, c, f, g**, **Supplementary Fig. 14, 15, 17, Video 4, 5**). We also proved that such direct 2D-3D VCD transformation is far superior to further post-enhancing 3D LFD results using established deep-learning restoration methods^19^ (**Supplementary Note Fig. 3**). Beyond the known limitation on recovery of dense signals using previous LFDM implementations, we further demonstrated that our VCD-LFM could reconstruct a 3D beating heart expressing highly-dense trabecular myocardium (*Tg(cmlc2:gfp)*), **Fig. 3h, i, Supplementary Fig. 16, Video 6**), and the limit was relatively high (**Supplementary Fig. 18**). Furthermore, the VCD-Net also generalized well when trained on one type of cardiac sample and applied to another, or trained on hybrid cardiac datasets together and applied to all of them (**Supplementary Note Fig. 4.1, 4.2)**. The fast VCD-LFM visualization with accurate single-cell resolution (**Supplementary Fig. 19**) then permitted quantitative investigation of transient cardiac hemodynamics. We tracked 19 individual RBCs throughout the entire cardiac cycle of 415 ms, during which the blood flow was pumped in-and-out of the ventricle at a high speed of over 3000 μm s^−1^ (**Fig. 3d, e**). With segmenting the continuous heart boundary, we successfully quantified the volume change of myocardium during the diastole and systole, and calculated the ejection fraction of the heartbeat (**Fig. 3j, k**). Furthermore, through a combination of VCD-LFM with SPIM, we realized 3D visualization and quantitative analysis of the blood flow inside the static zebrafish vessels (*Tg(fli1:gfp; gata1a:dsRed)*), **Methods, Supplementary 20, Video 7, 8**).

**Figure 3.**
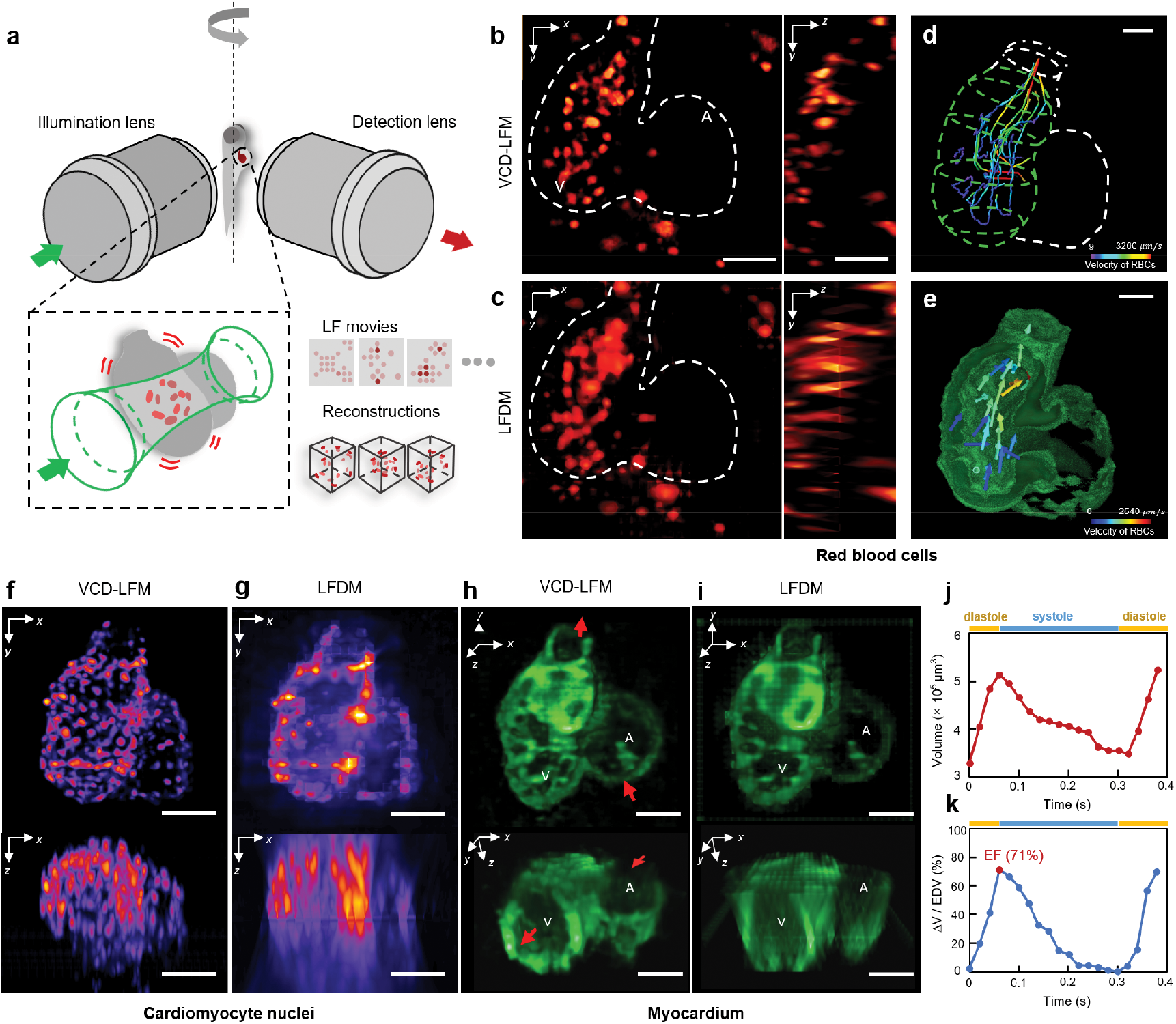
Imaging of various cardiac dynamics in beating zebrafish heart using VCD-LFM. (**a**) Schematic of selective volume illumination based light-field imaging for zebrafish experiments. High-contrast light-field sequences of flowing RBCs (beating heart) are recorded under a rod-like beam illumination setup that selectively excites the fluorescence signals within the volumes of interest. (**b**),(**c**) MIPs in *x-y* (left) and *y-z* (right) planes of one instantaneous volume of flowing RBCs by VCD-LFM and LFDM, respectively. The dash lines indicate the heart. (**d**) Tracks of 19 single RBCs throughout the cardiac cycle. A static heart is outlined for reference. (**e**) Velocity map of two temporally adjacent volumes of RBCs during systole. (**f**),(**g**) MIPs in *x-y* (top) and *x-z* (bottom) planes of one instantaneous volume of beating cardiomyocyte nuclei by VCD-LFM and LFDM, respectively. (**h**),(**i**) 3D visualization of beating myocardium in a transient moment by VCD-LFM and LFDM, respectively. The myocardium was densely labeled by GFP which shows continuous trabecular structures. Arrows indicate the inlet and outlet of cardiac pumping. A: atrium; V: ventricle. (**j**) Volume of the myocardium during the diastole and systole in one cardiac cycle. (**k**) Volume change ratio of the ventricle during the diastole and systole in one cardiac cycle. The rate is calculated by (*V*_*time*_ - *ESV*) / *EDV*, where *V*_*time*_ is the time-varying volume of the ventricle during heartbeat and *ESV* and *EDV* represent the volumes at the end of systole and diastole, respectively. The ejection fraction of the heartbeat (EF), given by (*EDV* - *ESV*) / *EDV*, also shown as the peak in the curve, is calculated to be ~71%. Twenty out of a total of 120 time points (1 of each 6) are selected for the analysis during a ~400 ms cardiac cycle. Scale bars, 50 μm. The data shown are representative of *n* = 8, 4, 3 independent fish for blood cells, cardiomyocyte nuclei and myocardium imaging, respectively.

To record the neural activities of acting *C. elegans* at single-cell resolution, previous studies captured the worm in 2D or 3D at a frame/volume rate up to 50 Hz^7, 17, 20^. An additional infrared channel is often required to provide bright-field images of worm behaviors as a reference. Our VCD-LFM, which is capable of 100-Hz recording and 13.5-Hz reconstruction of fast 3D processes combined with network-based all-fluorescence analysis, offers a new pipeline for the study of neural activities and relevant worm behaviors in whole moving worms, without motion blur or processing delay. For cardiovascular imaging, recent scanning-based approaches for volumetrically imaging the beating heart and blood flow in zebrafish larvae^21, 22^ have required complicated optics, ultra-fast camera, and high-intensity excitation. In contrast, our method based on a relatively simple system and easy-spread computational procedure offers a compelling solution for investigating the dynamic properties and functions of the cardiovascular system with single-cell resolution and low phototoxicity. Therefore, VCD-LFM could be a valuable tool for studying dynamics on fast timescales, potentially benefiting a variety of applications such as neural signaling-regulated phenotype studies and medically relevant dysfunctions of the heart and blood transport system in model organisms.

## Discussion

In summary, we introduced a VCD-LFM approach and demonstrated its ability to image transient biological dynamics with improved spatial resolution, minimal reconstruction artefacts, and increased reconstruction throughput. These advances are crucial for sustained volumetric imaging of blood flow in beating zebrafish heart and neural activities in behaving *C. elegans* at high resolution. The network-based VCD computational model is also robust, versatile, and ready for widespread application. While VCD-LFM greatly improves reconstruction quality from one originally coupled with the optics system to one can be optimized via prior data training, it still requires the pre-acquisition of training images. We expect this will improve with the continued development of deep learning. Besides the combination with basic LFM setup, we note that the VCD-Net is also compatible with modified LFM modalities, such as a dual-objective setup^11^ or scanning light-sheet illumination^18^. Finally, we expect that VCD-LFM could potentially bring new insights for computational imaging techniques and allow us to further push the spatiotemporal resolution limits for the observation of biological dynamics.

## Acknowledgements

We thank Jau-Nian Chen and Yuan Dong at UCLA for their help on zebrafish experiments. We thank Yanyi Huang at Peking University and Jianbin Wang at Tsinghua University for their discussions and comments on the manuscript. We thank Le Xiao for his help on the code implementation. We acknowledge the David Geffen School of Medicine at UCLA, for providing the Zebrafish Core Facilities and lab space. This work was supported by the following grants: National Key R&D program of China (2017YFA0700501 to P.F.), National Natural Science Foundation of China (21874052 and 21927802 to P.F., 31671052 to S.G.), National Science Foundation of Hubei (2018CFA039 to S.G.), National Institute of Health (NIH, HL149808, HL118650, HL129727, HL111437 to T.K.H., K99 HL148493 to Y.D.), Department of Veterans Affairs (VA, I01 BX004356 to T.K.H.), and the Junior Thousand Talents Program of China (P.F. and S.G.). Confocal experiment was done at the Advanced Light Microscopy/Spectroscopy Laboratory and the Leica Microsystems Center of Excellence at the California Nano Systems Institute at UCLA with funding support from NIH Shared Instrumentation Grant (S10OD025017) and NSF Major Research Instrumentation grant (CHE-0722519).

## Author contributions

P.F., Z.W. and H.Z. conceived the idea. P.F., S.G. and T.K.H. oversaw the project. Z.W., L.Z., G.L., Y.Y. developed the optical setups. H.Z., Z.W. and L.Z. developed the programs. Z.W., L.Z., G.L., Y.D. and Y.L. conducted the experiments, processed the images. Z.W., L.Z., H.Z., L.X., M.Z., S.G., T.K.H. and P.F. analyzed the data and wrote the paper.

## Methods

### Epi-illumination LFM setup

An epi-fluorescence light-field setup was built on an upright microscope (BX51, Olympus). The light-field/wide-field detection paths were appended to the camera port of host microscope, with using a flip mirror to switch between two detection modes. A motorized z-stage (Z812B, Thorlabs) together with a water chamber were directly mounted onto the microscope stage (x-y), to three-dimensionally control the samples inside the chamber. A water immersion objective (LUMPlanFLN 40×/0.8w, Olympus) was used to collect the epifluorescence signals from samples. For recording the light field, a microlens array (MLA, APO-Q-P150-F3.5 (633), OKO Optics) was placed at the native image plane to collect the light-field signals. A 1:1 relay system (AF 60 mm 2.8D, Nikon) was used to conjugate the back focal plane of MLA with the camera sensor plane (Flash 4.0 V2, Hamamatsu). The light-field path was optionally extended to dual-channel detection by dividing after MLA and adding an extra camera sensor for the *C.elegans* experiments. See **Supplementary Fig. 1** for more details of Epi-LFM setup.

### Selective plane/volume illumination LFM setup

We also developed a LFM setup based on selective volume illumination. Two pairs of beam reducers combined with an adjustable iris were used to generate a scalable rod-like beam (473 or 532-nm), which was finally projected onto the sample through a 4× illumination objective (Plan Fluor 4×/0.13, Nikon) placed perpendicular to the detection path. It confined the fluorescence excitation within the heart region of zebrafish embryo, reducing the excessive emission from out of the volume of interest that could smear the desired signals. This selective volume illumination mode provided light-field image with less background noise and increased contrast^18^. For observing the dynamic process of blood flowing through vessels, we also integrated a standard SPIM channel (473-nm, 4-*μm*-thickness laser-sheet) to implement the 3D imaging of steady vessels. The light-rod/light-sheet illumination paths were aligned, providing double excitation to the sample from its dual sides. A water immersion objective (Fluor 20×/0.5w, Nikon) were used to collect the dual-color fluorescence signals from the RBCs/cardiomyocytes. A dichroic mirror split the dual-channel GFP (vessels)/DsRed (RBCs) signals for wide-field and light-field detection, respectively. The light-field detection here followed the same design used in the epi-illumination LFM. See **Supplementary Fig. 2** for more details of this hybrid setup.

### VCD light-field reconstruction Network

In a general convolutional neural network (CNN), a certain *N*th convolutional layer receives feature maps from the last *(N-1)*th layer, and generates new feature maps using different convolution kernels. The network finally produces a multi-channel output, in which each channel is a non-linear combination of the original input. It’s reasonable to share this concept with the synthetic refocusing in the light-field imaging, where each synthetic focal plane of the reconstructed volume can be interpreted as a superposition of different views extracted from the light-fields. Through cascaded layers, our model is expected to gradually transform the original angular/view information from the light-field raw image into depth features, eventually forming the conventional 3D image stack, and reconstructing the scene. In our implementation, the customized VCD-Net is based on a modified U-Net architecture that contains a downsampling path and a symmetric upsampling path^23^. Along both paths, each layer has three parameters: **n, f** and **s**, denoting the output channels number, the filter size of convolution kernel and the step size of the moving kernel, respectively, as specified in **Supplementary Fig. 3 and Supplementary Table 1.** The pixels from the input 2D light-field raw image (dimension: **a** × **b**, height × width) are first reformatted into a series of different views (dimension: **a/d** × **b/d** × **d**^**2**^, height × width × views) according to their relative positions to each lenslet. A subpixel up-scaling part further interpolates these views to dimension **a** × **b** × **d**^**2**^(height × width × views). Then in the first VCD layer, the initial transformation from “view” to “channel” is done by convolutions with all these views using different kernels of each channel, generating an output with dimension **a** × **b** × **n**, where **n** is the channel numbers. The following convolution layers keep combining old channels from previous layer and generating new ones to excavate the hidden features from input image. Local residual connections are integrated into the downsampling path to fully extract the hierarchical features. Finally, the last layer outputs a 3D image with the channel number **n** equal to the desired number of synthetic focal planes **c**, thereby finishing the transformation from “channel” to “depth” (dimension: **a** × **b** × **c,** height × width × depth).

For VCD-Net training, HR 3D reference images were acquired from confocal microscopy of static samples or synthetic data. Then a light-field projection based on wave optics model^13^ was applied to these HR references to generate corresponding synthetic 2D light-field images as inputs (**Supplementary Note 1**). The VCD-Net was gradually trained via iteratively minimizing the difference between its intermediate outputs and the HR references. With setting appropriate loss functions, such as the MSE (Mean Square Error) of the pixel intensity, the VCD-Net obtained optimized kernel parameters for each layer and efficiently converged to a well-trained status, at which the network can transform the synthetic light-field inputs back to 3D images. At inference stage, the trained VCD-Net directly infers a sequence of 3D images from an input light-field video, which contains many light-field frames recording the dynamic biological processes. The time consumption for VCD-Net procedure depends on the dataset size and computation resources. As a reference point, the VCD-Net converged after training on 4580 pairs of blood cells image patches (size: 176 × 176 × (51) pixels) with 110 epochs. The time cost was ~4 hours on a single GPU. Then the trained network spent ~15 seconds to reconstruct 450 consecutive volumes (size: 341 × 341 × 51 pixels) from acquired light-field videos. This 4D reconstruction throughput was compared to ~11.8 hours (~42467 s) by running LFD (8 iterations) on the same workstation. The computation was performed on a workstation equipped with Intel(R) Core (TM) i9-7900X CPU @3.3GHz, 128G RAM, and Nvidia GeForce RTX 2080 Ti graphic cards. For more details in VCD-Net training and inference, see **Supplementary Note 2** and our open-source code. The comparative performance of our direct 2D-3D VCD-Net recovery and LFD recovery (2D-3D) plus other deep-learning image restoration (3D-better 3D) was shown in **Supplementary Note 3**. The generalization ability of VCD-Net, reflected by the hybrid data training/multi-sample recovery and cross-sample/transfer learning application, was discussed in **Supplementary Note 4**.

### PSF measurements

We used both light-field (40×/0.8w objective) and 3D wide-field (40×/0.8w objective plus 0.5-μm z-step) modes to image the same volume of sub-diffraction fluorescent beads (0.5 μm Lumisphere, BaseLine) distributed in a piece of hydrogel (0.7% low melting agarose solution, BBI Life Sciences). To demonstrate the tunable performance of VCD-Net with applying different training data, we trained anisotropic and isotropic VCD-Nets using synthetic anisotropic and isotropic beads, respectively. These synthetic beads with random 3D distributions were generated using a 3D Gaussian kernel with controllable FWHMs (anisotropic: 1 × 1 × 3 μm; isotropic: 1 × 1 × 1 μm). Then 3D image stacks with 60-μm depth were recovered from the recorded light-field image using LFD, anisotropic and isotropic VCD-Nets. The reconstruction yielded 61 z planes with voxel size 0.34 × 0.34 × 1 μm for anisotropic VCD-Net and LFD (8 iterations), and 177 z planes, with voxel size 0.34 × 0.34 × 0.34 μm for isotropic VCD-Net. The reconstructed beads were then detected and fitted with 1D Gaussian function in each dimension to determine the full width at half maximum (FWHM) as spatial resolution metrics using a custom-written MATLAB script. The PSFs of LFD and VCD-Nets at certain depth were then compared via plotting the line profiles of the same resolved beads. The achieved lateral and axial resolutions at certain depth by each method were indicated through calculating the averaged FWHM values of the resolved beads, e.g., z = −14 μm. The performance of anisotropic and isotropic VCD-Net *versus* LFD at different depths were further analyzed via measuring the FWHMs of resolved beads across a 60-μm depth, with their variation indicating the non-uniformity of light-field recovery.

### *C. elegans* strain

The strain ZM9128 *hpIs595[Pacr-2(s)::GCaMP6(f)::wCherry]*, with calcium indicators tagged to the A-and B-class motor neurons, was used to detect the neural activities in the acting worm (**Fig. 2**). The strain QW1217 *hpIs467[Prab-3::NLS::RFP]*, with nuclei labelled in all the neuron cells, was used to demonstrate the imaging performance of VCD-LFM (**Supplementary Fig. 12**). All *C. elegans* were cultured on standard Nematode Growth Medium (NGM) plates seeded with OP50 and maintained at 22 °C incubators until L4 stage.

### *C. elegans* imaging

To obtain HR 3D data for network training, the L4-stage anesthetized worms (QW1217 *hpIs467*, ZM9128 *hpIs595*, by 2.5 mM levamisole in M9 buffer, Sigma-Aldrich) were first imaged using a 40×/0.95 objective on a confocal microscope (FV3000, Olympus). To demonstrate the performance of VCD-LFM, anesthetized worms (QW1217 *hpIs467*) embedded in agarose were *in situ* imaged by both light-field and wide-field detection modes in our Epi-illumination LFM. The acquired 3D wide-field image stacks (1-μm step size) were deconvolved in Amira software (Thermo Scientific), for comparison with those light-field reconstructions by LFD and VCD-Net. The 3D reconstructions encompassed 31 z planes, with voxel size of 0.34 × 0.34 × 1 μm. In the behavior studies of *C. elegans*, the awake L4-stage worms (ZM9128 *hpIs595*) were loaded into a microfluidic chamber (~300 × 300 × 50 μm), which allows the worms to act within the FOV of a 40× objective. The GCaMP/RFP signals of acting worm were then recorded for one minute at 100 Hz frame rate (2-ms exposure time).

### Quantitative Analysis of neural activities and behavior of acting *C. elegans*

We performed semi-automated tracking of the movement and intensity fluctuation of each individual neuron using TrackMate Fiji Plugin^24^. Neurons were detected automatically in each volume with applying a circular ROI through the Difference of Gaussian (DoG) detector and then tracked using Kalman filter. In cases when this automatic tracking failed due to fast movement of the neuron, the missing detections and tracking mistakes were manually corrected. For each neuron in each volume of GCaMP or RFP channel, all the pixels within the ROI of certain neuron were averaged to generate a single value *F*_*g*_ or *F*_*r*_ representing the fluorescence intensity of this neuron in each channel. To extract calcium dynamics, we measured *(F* − *F*_O_)/*F*_O_, where *F* = *F*_*g*_ / *F*_*r*_, also known as ratiometric correction, is the ratio of GCaMP fluorescence *F*_*g*_ to RFP fluorescence *F*_*r*_, and *F*_O_ is the neuron-specific ratiometric baseline being the average of the lowest 100 values. Worm behavior analysis was then implemented based on the same fluorescence images. We developed an efficient worm analysis pipeline which could 1. rapidly infer the worm outlines throughout the whole time period using a U-Net based image segmentation^23^; 2. extract the center lines from the segmented outlines to calculate the changing curvatures and motion velocities^25^. More details are given in **Supplementary Note 5**. The body curvature map shown in Fig. 2f indicates the time-varying worm postures. The velocity curve shown in Fig. 2g represents the displacement along the worm body direction.

### Fish husbandry and lines

Transgenic zebrafish lines *Tg(gata1a:dsRed; cmlc2:gfp)* (**Fig. 3**, **Supplementary Fig. 14, Video 4, 9**), Transgenic zebrafish lines *Tg(cmlc2:gfp)* (**Fig. 3**, **Supplementary Fig. 16, Video 6**), *Tg(fli1:gfp; gata1a:dsRed)* (**Supplementary Fig. 14, 20, Video 7, 8**), *Tg(myl7:nls-gfp)* (**Supplementary Fig. 15, Video 5**) were used in our experiments. Embryonic stage fishes were maintained until 3-4 dpf in standard E3 medium, which was supplemented with extra PTU (Sigma Aldrich, MO) to inhibit melanogenesis. Then, the larvae were anesthetized with tricaine (3-amino benzoic acidethylester, Sigma Aldrich, MO) and immobilized in 1% low-melting-point agarose inside FEP (Fluorinated Ethylene Propylene) tube for further imaging. In the imaging of cardiac blood flow, the embryos were injected with gata2a morpholino oligonucleotide at single-cell stage, to slow down hematopoiesis, and thereby reduce the density of RBCs. All the experiments were performed in compliance with UCLA IACUC protocols.

### Zebrafish imaging

The fish samples were precisely moved by a custom stage (xyz + rotation) so that their signals of interest could be positioned within the optimal imaging region, determined by the size of selective volume illumination and the FOV of detection objective (LUMPlanFLN 20×/0.5w, Olympus). During cardiac imaging, the middle of heart was moved to the focal plane so that the volumetric illumination (473 or 532-nm) could selectively excite the RBCs/myocytes nuclei/myocardium. The light-field movies were recorded at 200 Hz frame rate, with 768 × 768 frame size that corresponds to a lateral FOV of ~250 × 250 μm. The movies covered 4-5 cardiac cycles with containing 450 frames (5 ms exposure for each frame). For imaging the dynamic process of blood flow inside vessels, the cameras of dsRed and GFP channels recorded light-field video of rapidly flowing RBCs and SPIM image stack of static tail vessels, respectively. The light-field movies contained 600 frames for each one (10 or 5 ms exposure). The SPIM stack contained 50 planes that covered a 100-μm z depth of fish tail (step size 2 μm).

To obtain HR 3D images of RBCs, myocyte nuclei and myocardium for VCD-Net training, deeply anesthetized fish larvae with immobilized hearts were embedded in 1% low-melting agarose for sustained confocal imaging (SP8-STED/FLIM/FCS, Leica) using a 20×/0.75 Objective (HC PL APO CS2). Then we trained three VCD-Nets based on these HR images of static blood cells (16 fishes), nuclei (23 fishes), and myocardium (11 fishes), and their paired LFPs. Besides the confocal imaging of immobilized heart, we also combined SPIM with retrospective gating method^21, 26, 27^, to obtain the light-sheet based HR 3D image of periodically beating myocardium for network training. A movie stack of light-sheet images of the dense myocardium was first acquired, containing 60-70 movies at different planes (z step = 2 μm) that captured the complete myocardial structures. Each movie was then temporally aligned using the retrospective gating, to reconstruct a beating 3D heart. When the training using these prior data was finished, the network inferred 3D images from empirical light-fields with output voxel size 0.68 × 0.68 × 2 μm (blood cell, myocardium) or 0.326 × 0.326 × 3 μm (myocyte nucleus). The output image stacks contain 51 planes with depth spanning from −50 to 50 μm or −75 to 75 μm.

### Resolution quantification of VCD-Net and LFD reconstructions

We applied decorrelation analysis^28^ to quantify the resolutions of VCD-Net and LFD reconstructions. The analysis was performed using ImageJ plugin (image decorrelation analysis). We set recommended parameters (Radius: 0-1; Nr: 50; Ng:10) to calculate the cut-off frequency *k*_*c*_, which represents the derivative of the smallest resolvable distance in the reconstructed images. **Supplementary Fig. 15-17** show the higher *k*_*c*_ obtained in VCD-Net results than LFD results, which quantitatively validate the better resolution achieved by VCD-Net. We also used the Fourier spectrograms (by ImageJ plugin), to further confirm the resolution advantage of VCD-Net.

### Velocity map of RBCs and volume-based ejection fraction analysis of myocardium in zebrafish heart

Tracking of the flowing RBCs was applied using Imaris (Bitplane). **Fig. 3d** shows the trajectories of 19 RBCs throughout one cardiac cycle (415 ms). **Fig. 3e** demonstrates the velocity map that has been extracted at one specific time point during systole, which contains 31 vectors from all trackable RBCs in that frame. The analysis was written in Matlab and the vector field was visualized by Mayavi^29^. Volumetric segmentation of beating myocardium was performed using Amira. A blow tool was used to allow semiautomatic definition of the inner boundary of the ventricle at each slice. Once all the slices in certain 3D image stack of one time point were correctly segmented, the software could calculate the volume of segmented ventricle based on the slice thickness and the defined area. After the volume of the beating heart at all time points being calculated, we obtained the volume change ratio of the ventricle during the diastole and systole in one cardiac cycle.

### Reporting Summary

Further information on research design is available in the Nature Research Reporting Summary linked to this article.

## Data availability

The datasets generated and analyzed in this study are available from the corresponding author upon request. Example model checkpoints and data are packaged with the code for usage tutorial. Source data for main figures are provided with this paper.

## Code availability

Custom codes for VCD-Net and quantitative analyses implemented in current study are either available at https://github.com/feilab-hust/VCD-Net or from the corresponding authors upon request.

