## Supplementary Figures and notes for "View-channel-depth light-field microscopy: real-time volumetric reconstruction of biological dynamics by deep learning"

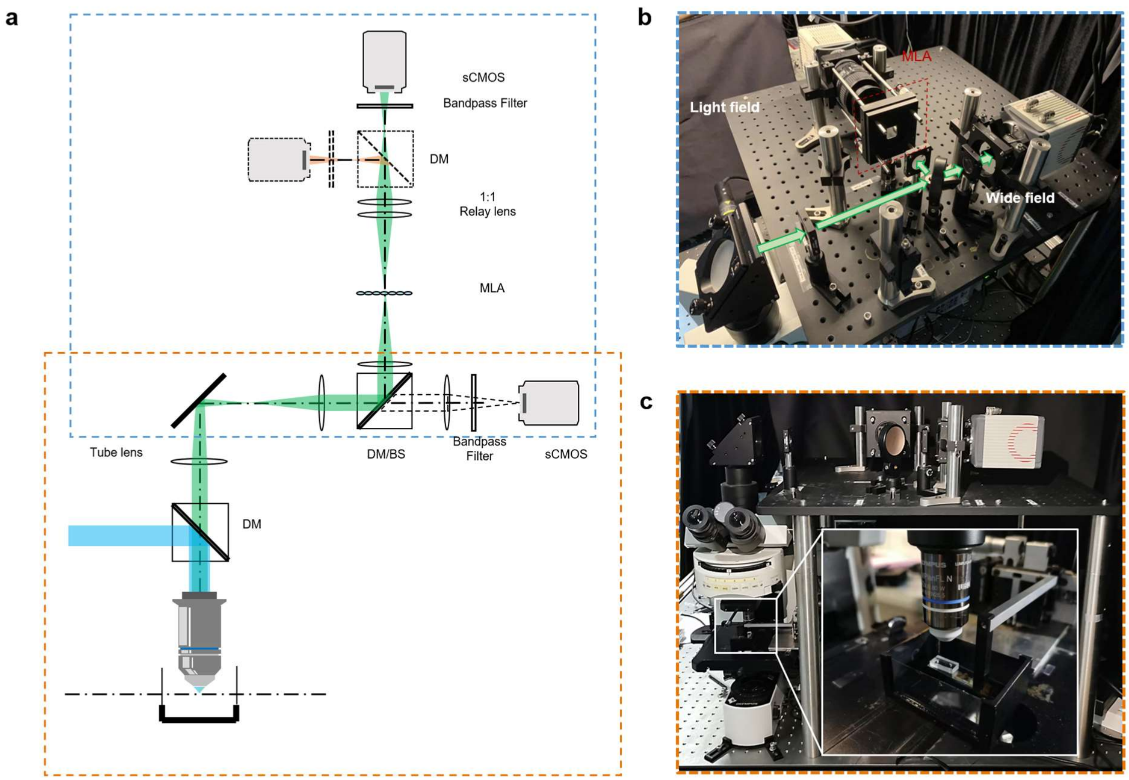


**Supplementary Figure 1**

**Epi-illumination light-field/wide-field microscopy setup.**

(**a**) Schematic drawing of our epi-illumination light-field/wide-field setup for beads and *C. elegans* experiments. The fluorescence signals collected by the detection objective (LUMPlanFLN 40×/0.8w NA, Olympus) were either focused onto the camera sensor 1 (Flash 4.0 V2, Hamamatsu) for wide-field imaging or onto the microlens array (APO-Q-P150-F3.5 (633), OKO Optics) for light-field imaging. In the latter case, a 1:1 relay system (AF 60 mm 2.8D, Nikon) was used to focus the camera sensor 2 (Flash 4.0 V2, Hamamatsu) on the back focal plane of MLA. DM (Dichromatic mirror) or BS (Beam splitter) was used for sequential or simultaneous light-field/wide-field imaging, respectively, depending on the experiments. A DM can be optionally added in front of the sCMOS in the light field path for dual-color light-field imaging. The system was based on an epifluorescence upright microscope (Olympus, BX51), as shown in (**c**), with its customized wide-field/light-field detection paths shown as (**b**). A microfluidic chip (inserted picture) was used to permit the worm acting within the FOV of 40× objective.

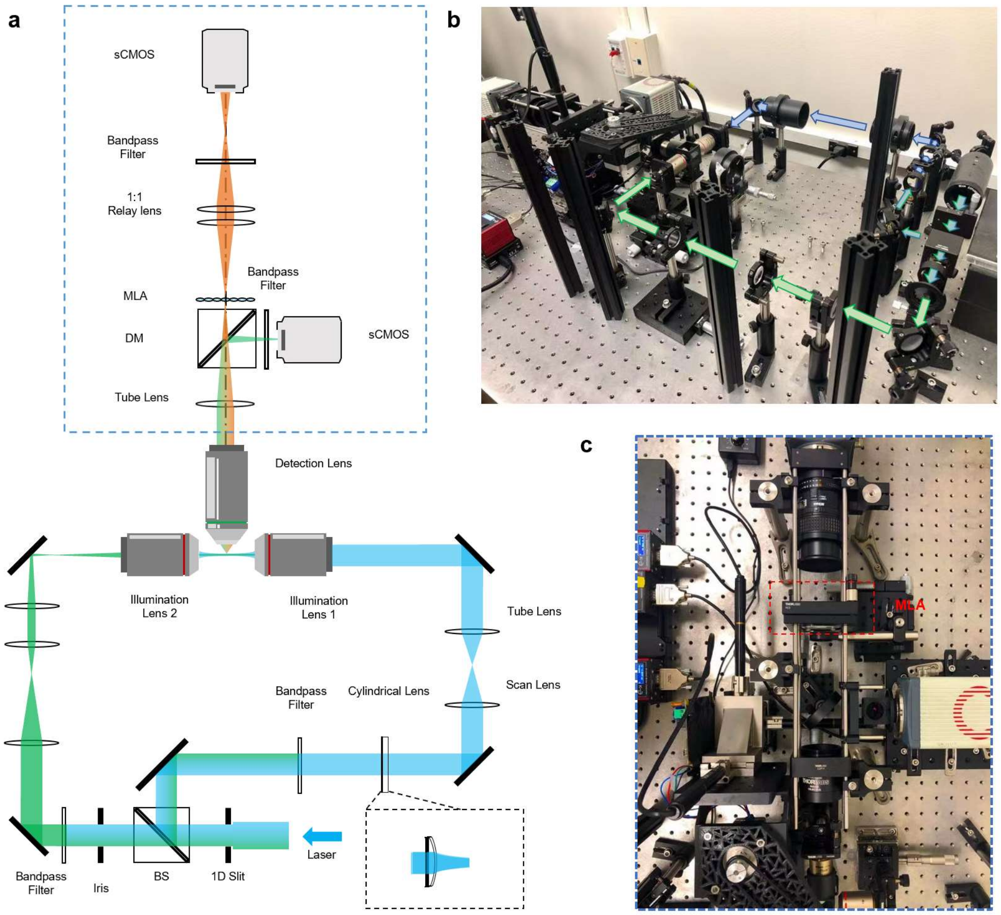


**Supplementary Figure 2**

**Selective plane/volume illumination based LFM setup.**

(**a**) Schematic drawing of our self-built selective plane/volume illumination setup for hybrid SPIM/LFM imaging of hemodynamics. A tunable rod-like laser beam (100-200 μm) for volumetric excitation and a thin laser-sheet (~4 μm) for planar excitation could be projected to the sample simultaneously, thereby enabling light-field and light-sheet imaging, respectively. (**b**) Overview of the system. (**c**) Top view of the dual-mode light-field/wide-field detection.

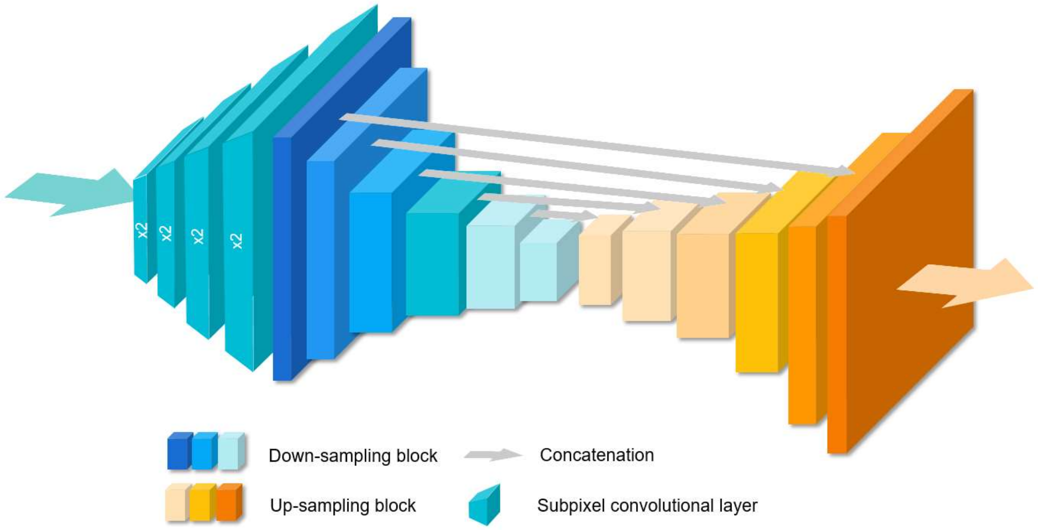


**Supplementary Figure 3**

**VCD-Net structure overview.**

Our approach adopted the well-known U-Net[1] for feature extraction and image reconstruction. Before the U-Net part, we designed a sub net to interpolate the extracted views into its original size (i.e., the size of the light-field measurement). The interpolation net contains 4 subpixel convolutional layers[2], with each up-scaling the image by a factor 2. The following U-Net has down-sampling and up-sampling parts for features encoding and decoding, respectively. The down-sampling part contains down-sampling blocks for feature extraction, where each down-sampling block consists of a convolutional layer, an activation function, a batch normalization and a max pooling. Accordingly, up-sampling blocks in the up-sampling part up-scales the feature maps. Each up-sampling block consists of a convolutional layer, an activation, a batch normalization and an up-sampling operation. Concatenations merge the feature maps of the corresponding down-sampling and up-sampling blocks. Detailed parameters of each layer are listed in **Supplementary Table 1**.

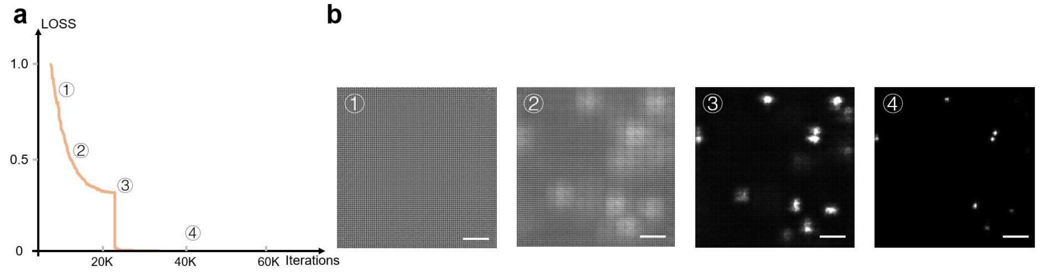


**Supplementary Figure 4**

**Iterative convergence of the network.**

(**a**) The decreasing tendency of the loss function (MSE between the outputs and the targets) during the network training. The parameters of the neural networks were initialized randomly. The training process updated them iteratively so that the network outputs would be more and more similar to the targets, with the decrease of the value of the loss function. (**b**) The intermediate outputs of the VCD-Net at different training time points shown in **a**. Scale bar: 10 μm.

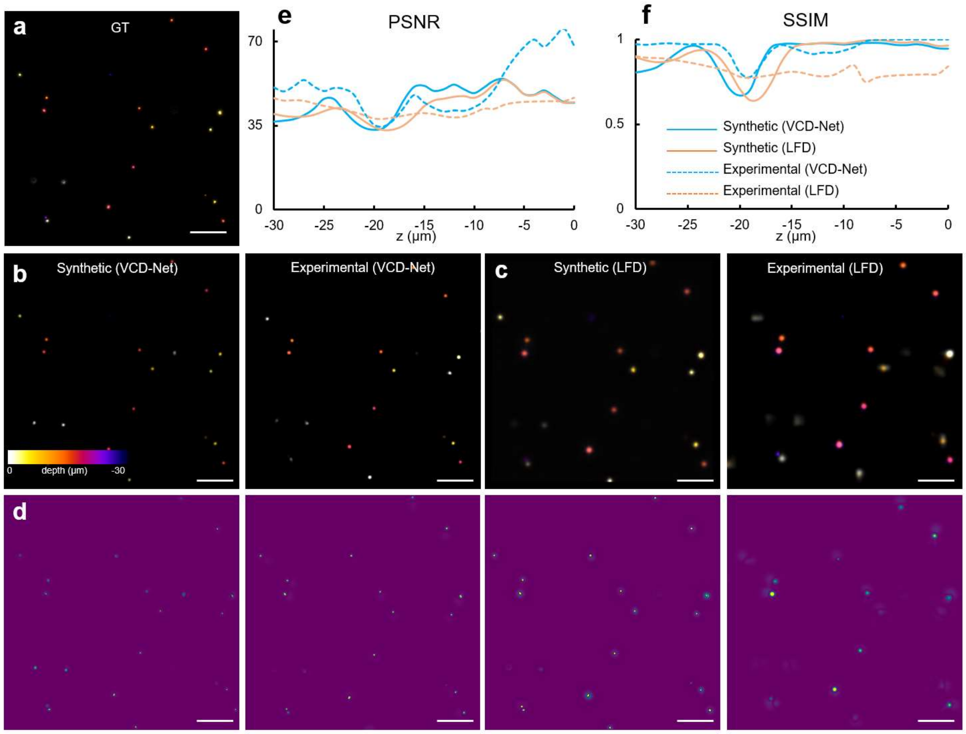


**Supplementary Figure 5**

**Comparison of reconstruction quality by VCD-Net and LFD.**

A 30-μm depth volume was *in-situ* imaged using light-sheet and light-field modes by selective plane/volume illumination LFM setup. As a result, we obtained HR ground truth (GT) by SPIM (**a**), experimental light-field image by LFM, and synthetic light-field image via the LFP of GT. Then we reconstructed 3D images from the experimental and synthetic light-field images using both VCD-Net and LFD, as shown in (**b**) and (**c**) respectively. Accordingly, their reconstruction fidelity could be intuitively compared through generating their error maps with GT, shown as (**d**). Furthermore, we quantitatively compared the reconstruction fidelity of VCD-Net and LFD via layer-by-layer calculating their peak signal to noise ratio (PSNR) and structure similarity (SSIM) indices, with using GT as reference. The results are shown in (**e**) and (**f**), respectively. Scale bar: 10 μm.

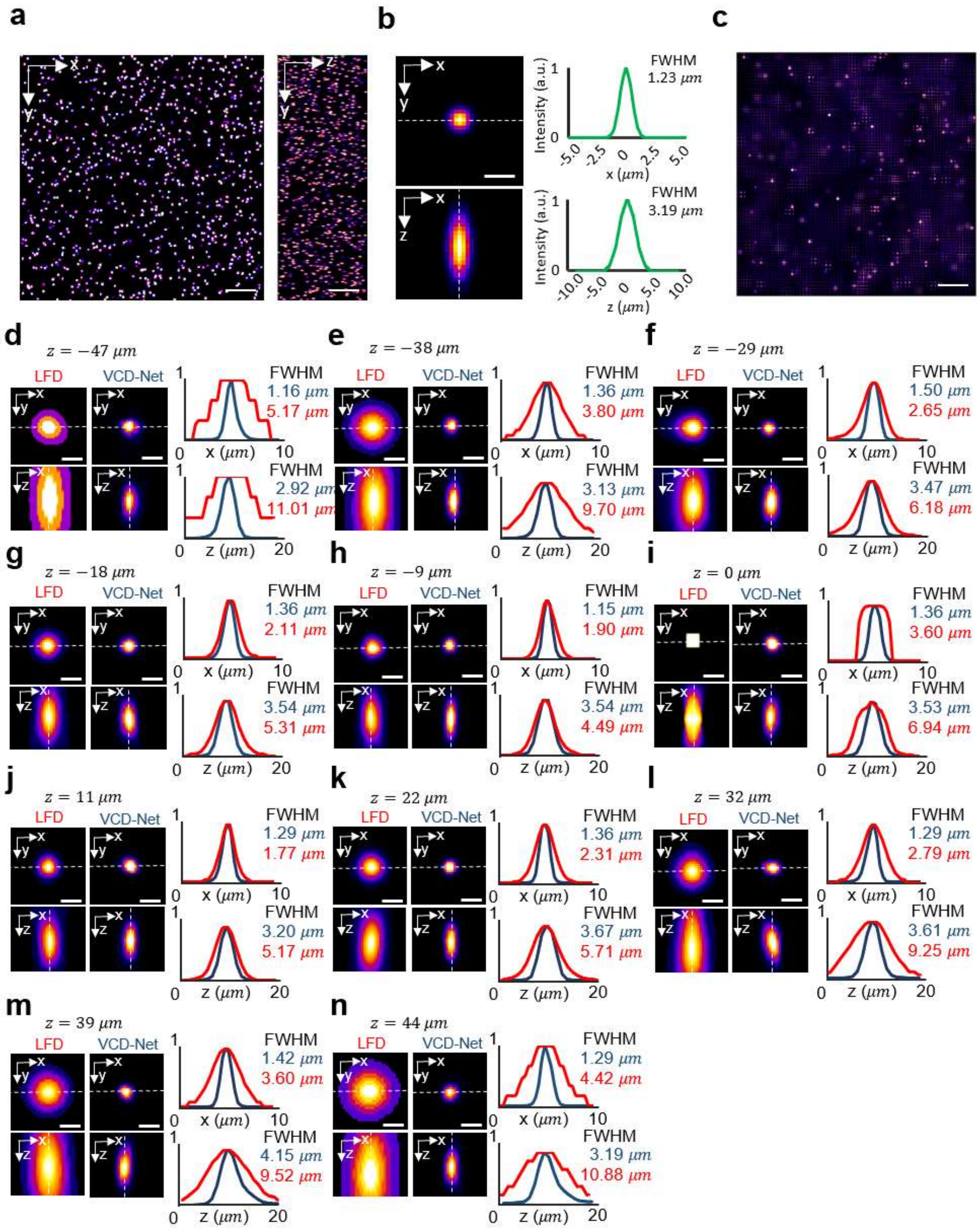


**Supplementary Figure 6**

**Comparison of full width at half maximum (FWHM) of beads at various depths reconstructed by LFD and VCD-Net.**

(**a**) Maximum Intensity Projection (MIP) of synthetic 3D beads. Beads were randomly distributed in in a volume of ~ 300 μm × 300 μm × 120 μm. Scale bar, 40 μm. (**b**) Individual reconstructed bead in (**a**) in lateral(x-y) and axial(x-z) dimensions, and the corresponding profiles along the dashed lines. Scale bar, 3 μm. (**c**) Synthetic light-field raw image of (**a**). Scale bar, 40 μm. (**d**)-(**n**) Cross-section images of reconstructed beads using LFD with 8 iterations (left column, red) and VCD-Net (right column, blue) at depths ranging from -47 μm to 44 μm. The profiles along dashed lines were plotted for each depth along lateral and axial directions. Scale bar, 3 μm.

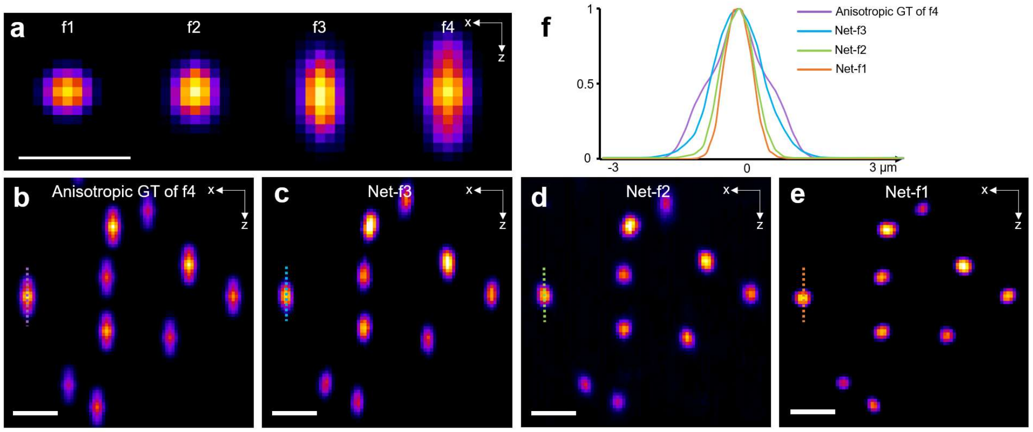


**Supplementary Figure 7**

**Training data oriented isotropic VCD reconstruction.**

We demonstrated that decoupled with the optical limitation, the reconstruction of VCD-Net could be optimized by applying higher-quality training dataset, and capable of reaching a near-isotropic resolution with isotropic training data. (**a**) Maximum intensity projections (MIPs) of the synthetic beads with various axial prolongation being 1 (f1, isotropic) to 4 (f4, anisotropic) times larger than the lateral width. (**b**) MIP of f4 anisotropic image stack. A synthetic light-field image was generated from it and used as the input for the following VCD-Nets test. (**c**)-(**e**) MIPs of reconstruction results by VCD-Nets. They have been trained on f3, f2 and f1 dataset respectively. (**f**) Line profiles of the resolved beads shown in (**b**) to (**e**). It's shown that by applying higher quality data training, the VCD-Net is capable of delivering data-guided reconstruction, with potential isotropic resolution beyond that of the source 3D image. Scale bar, 5 µm.

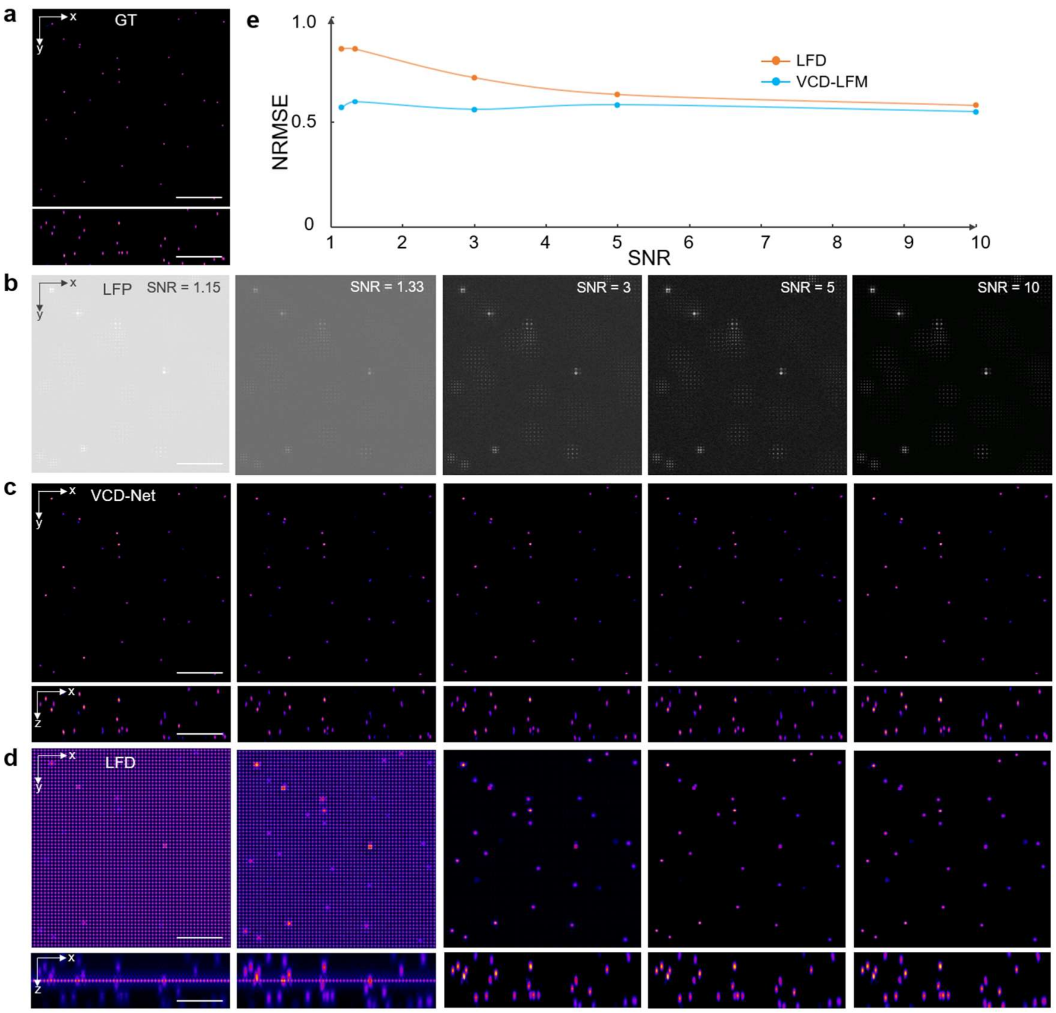


**Supplementary Figure 8**

**Comparison of VCD-Net and LFD on recovering low-SNR noisy signals.**

(**a**) MIPs in xy and xz planes of high-resolution high-SNR 3D image by light-sheet microscopy. (**b**) Five synthetic light-field images of (**a**) with different levels of background noise (Gaussian noise at LFP and Poisson noise at final sensor image) added. The calculated SNR is from 1.15 to 10. (**c**)-(**d**) MIPs in xy and xz planes of the 3D reconstructions of these different-SNR light-field images by VCD-Net and LFD, respectively. The VCD-Net was trained with different SNR light-fields with ground truth high-SNR 3D images. (**e**) Reconstruction error, termed normalized root mean square error (NRMSE, lower is better), of VCD-Net and LFD under different SNR conditions. Results in (**c**) to (**e**) validate the robustness of VCD-Net for recovering noisy data. Scale bars, 50 µm.

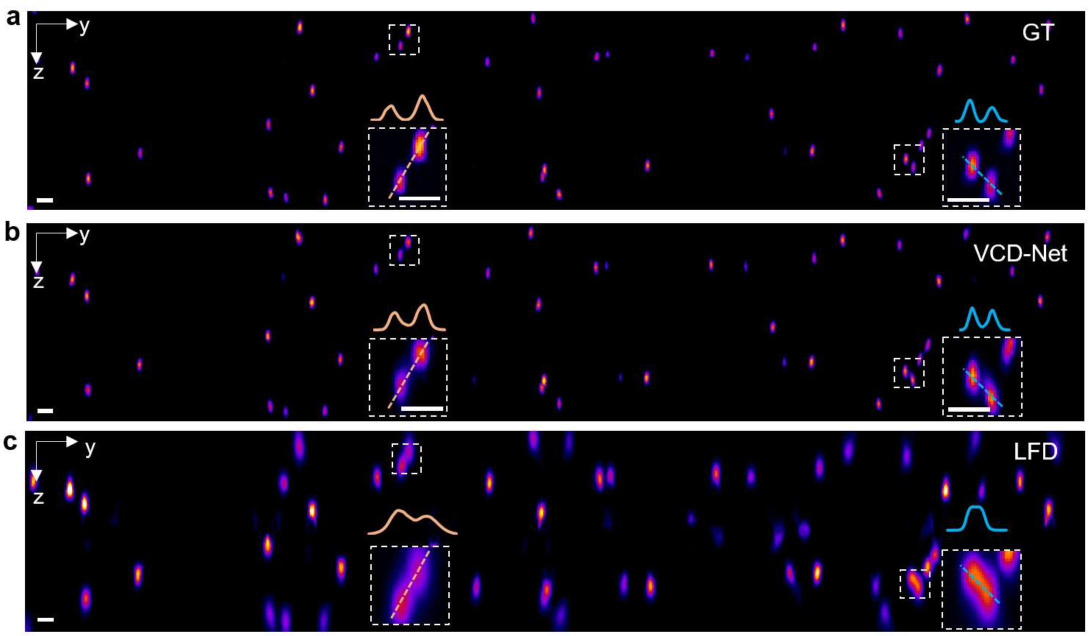


**Supplementary Figure 9**

**Performance of VCD-Net and LFD on resolving adjacent point signals.**

(**a**) High-resolution cross-sectioning image (y-z plane) of fluorescent beads by 3D wide-field microscopy (GT). (**b**),(**c**) 3D light-field reconstructions from the light field measurement of the same beads by VCD-Net and LFD, respectively. The comparative magnified views as well as the intensity profiles (along dash lines) have verified that while LFD failed to discern those closely adjacent points, VCD-Net clearly resolved them with accurate intensity and location shown. Scale bars, 5 μm.

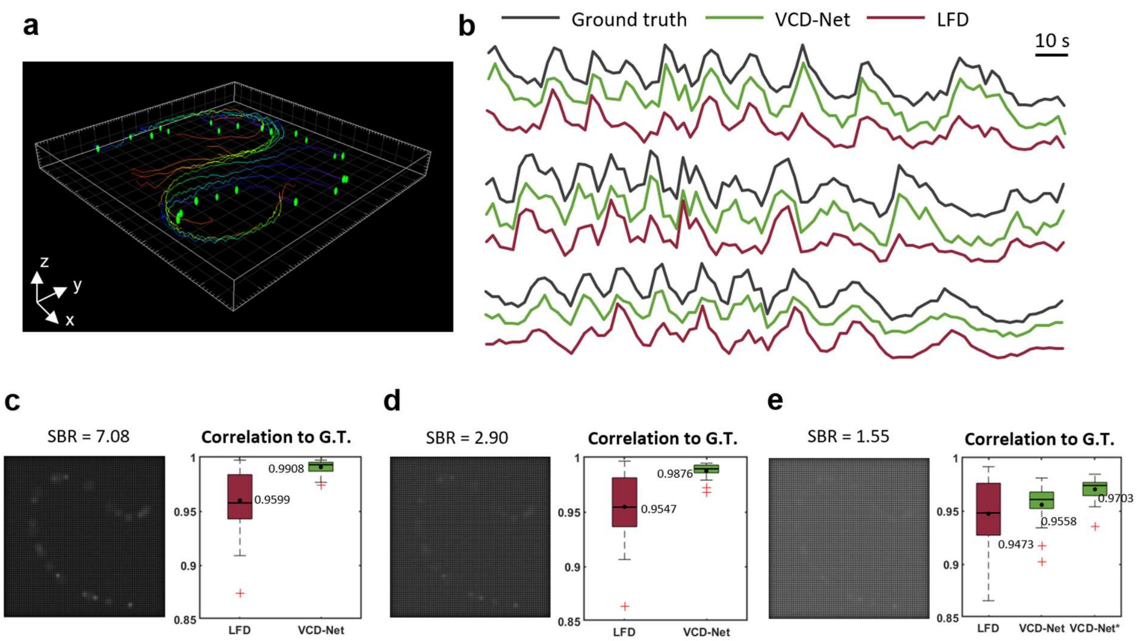


**Supplementary Figure 10**

**VCD-Net performance on tracking the signal fluctuation in moving neuron activities.**

We generated sequence of volumes containing synthetic point-like neurons with time-varying positions and intensities, to simulate the ground truth status of a GCaMP-labelled free moving worm. The positions were collected from empirical data (**Supplementary Video 2**) and calcium signals adopt the measurements in published work[3]. The synthetic light-field movie was generated and then input into the trained VCD-Net to reconstruct a 3D movie. **(a)** Trajectories of neurons (neuron candidate in green dot; trajectory color indicates the time: purple, beginning; red, end) within the first 5 seconds. **(b)** Intensity fluctuations of three selected signals, which simulated the calcium signaling of VA10, VA11 and DA7 neurons of the worm, in ground truth (black lines, upper), VCD-Net (green lines, middle) and LFD (red lines, lower) reconstructions, were compared throughout a 180-second period. **(c),(d),(e)** Comparative analysis correlating the ground-truth signals with VCD-Net and LFD signals reconstructed from light-field images at different signal-to-background ratios (SBR, 7.08 to 1.55), with consideration to samples with strong baseline background and noise. Gaussian and Poisson noise were added while the neural signal magnitude was altered to achieve different SBRs. Boxplots show the median as thick lines, and the 25th and 75th percentiles as box edges. Whisker: maximum and minimum after excluding outliers (red cross). Dot and number, the mean correlation. The VCD-Net was trained on high-SBR data and applied to all the three different SBR levels. VCD-Net* specifically trained on low-SBR data further improved the accuracy. The signal traces shown in **(b)** were based on the results in **(e)**.

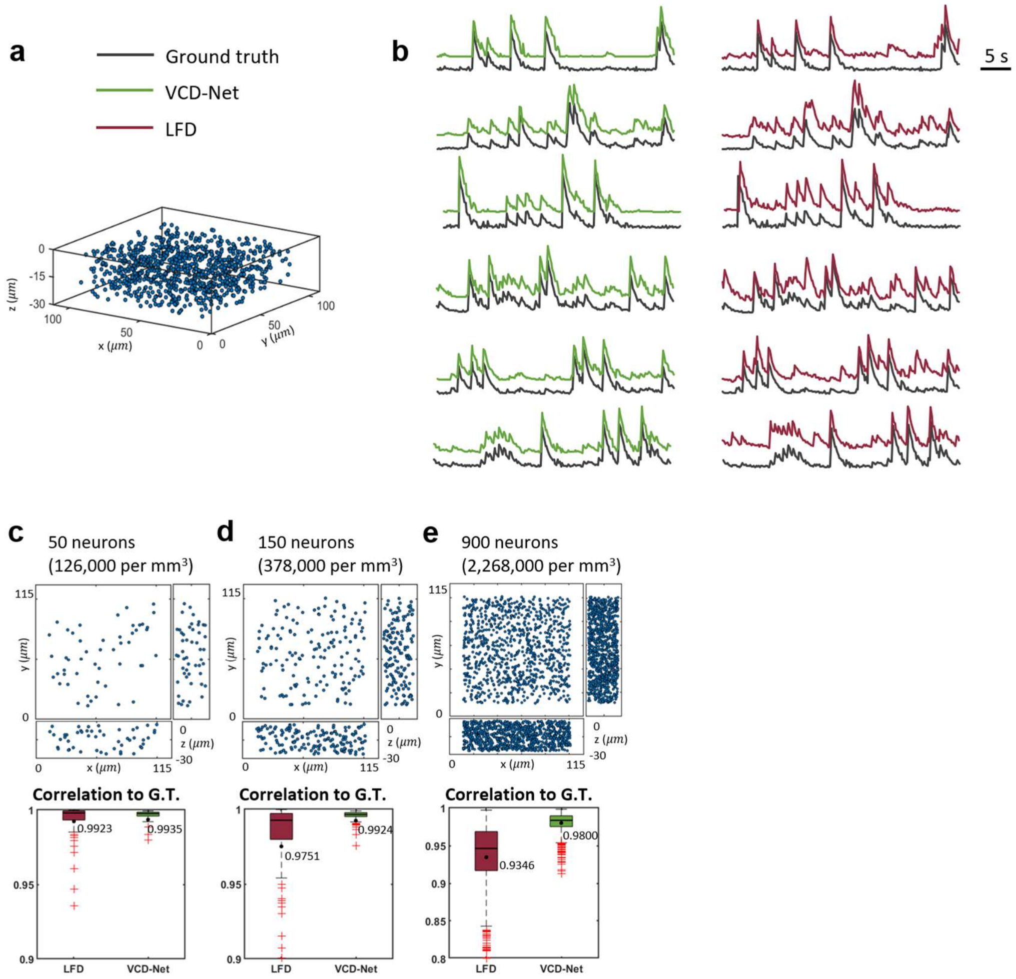


**Supplementary Figure 11**

**VCD-Net performance on static neuron activities with different signal densities.**

**(a)** Synthetic firing neurons with fixed positions and various densities were generated. Neurons (spheres of $8\ \mu m$ diameter) were randomly seeded in the volume of $116 \times 116 \times 30\ \mu m^3$ with signals simulated as Poissonian spike trains (firing rate 0.1 Hz, 200 time step at sampling rate 5 Hz) convolved with an exponentially decaying kernel (mean decay time constant 1.2s). **(b)** After reconstruction by VCD-Net (green lines, left panel) and LFD (red lines, right panel), neuron signals have been measured to compare with the Ground truth (black lines). 6 example traces are shown. **(c),(d),(e)** Intensity accuracy of reconstruction at different neuron densities. Boxplots show the median as thick lines, and the 25th and 75th percentiles as box edges. Whisker: maximum and minimum after excluding outliers (red cross). Dot and number, the mean correlation. The signal intensity traces shown in **(b)** were based on the results in **(e)**.

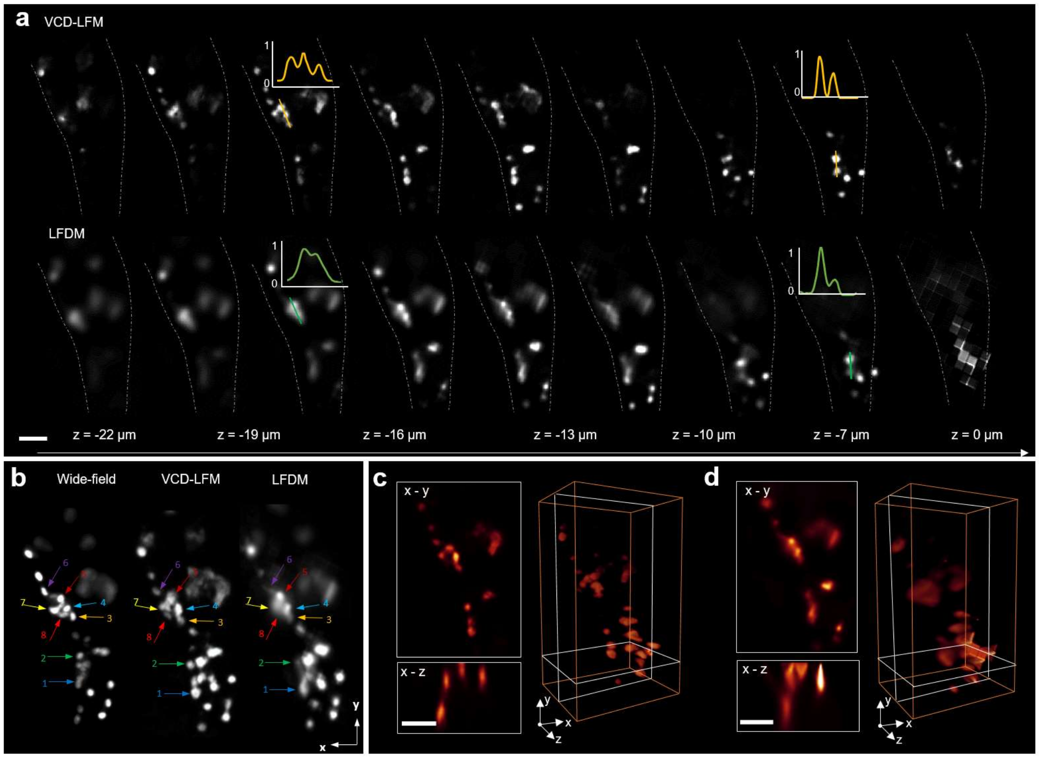


**Supplementary Figure 12**

**VCD-LFM, LFDM and 3D wide-field imaging of C. elegans.**

(**a**) Volumetric reconstruction of *C. elegans* (strain QW1217 *hpIs467[Prab-3::NLS::RFP]*) by our VCD-Net and LFD. The LFD result was generated after 8 iterations. Intensity profiles of the color lines at two planes (-18 µm and -7 µm) are drawn for each method. (**b**) MIP of reconstructed worm, for a clear illustration of ability of our VCD-Net to discern dense signals at the worm tail. The results were compared to the 3D wide-field results. Arrows and numbers indicated the neurons, which were successfully resolved in our VCD-Net result (middle) but blurred in LFD result (right). (**c**),(**d**) 3D renderings and cross sections of worm's tail reconstructed by our VCD-Net (left) and LFD (right), respectively. Scale bar, 10 µm.

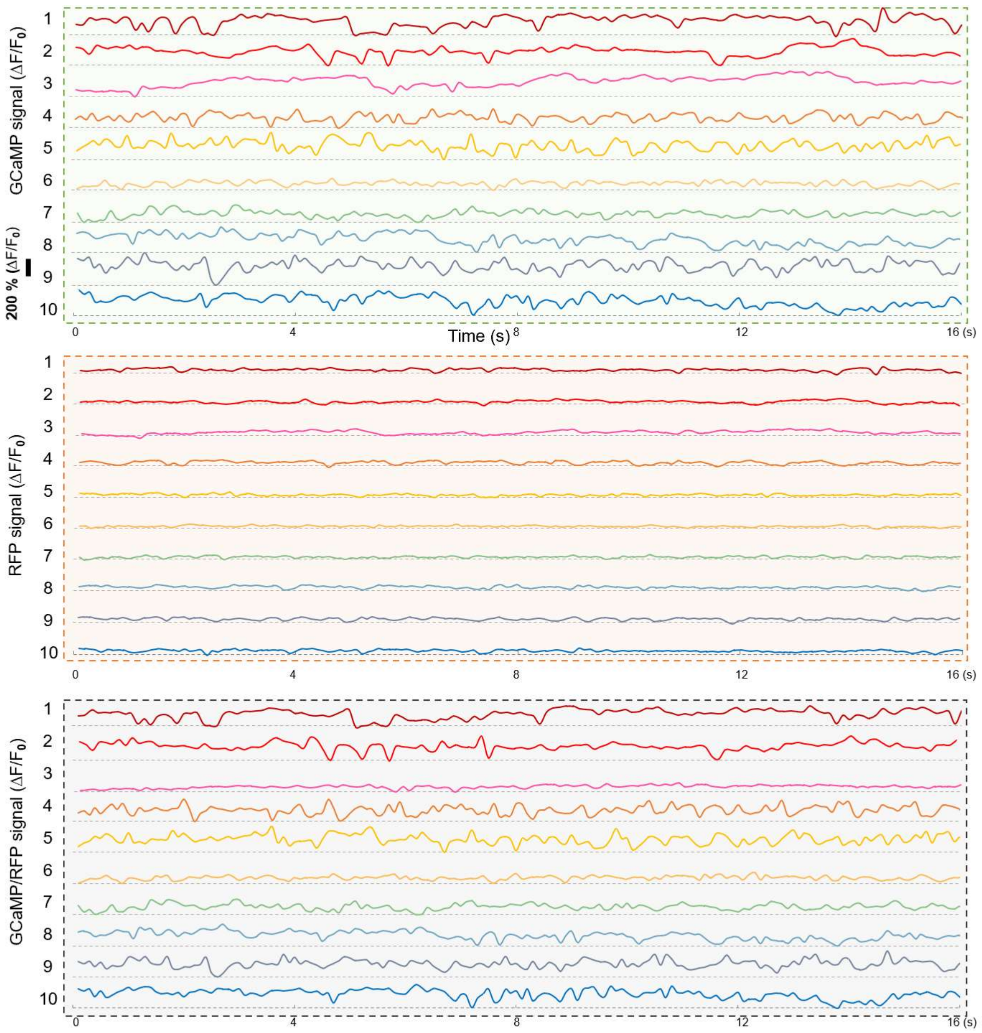


**Supplementary Figure 13**

**Dual-channel imaging of neural activities in freely moving C. elegans using VCD-LFM.**

L4-stage *C.elegans* with motor neurons labelled by GCaMP and RFP fluorescence (strain ZM9128 *hpIs595[Pacr-2(s)::GCaMP6(f)::wCherry]*) were imaged and reconstructed by dual-channel VCD-LFM. The fluorescence intensity fluctuations of GCaMP, RFP, and GCaMP / RFP signals in ten neurons are shown over a 16-second time period. The traces show relatively small intensity fluctuation in RFP signals as compared to GCaMP signals. Meanwhile, the ratiometrically-corrected GCaMP / RFP signals are still significantly correlated to the GCaMP signals after the motion artifacts being cancelled out. $F_0$ is the average of the lowest intensity values, and $\Delta F$ denotes the subtraction of current intensity $F(t)$ with $F_0$.

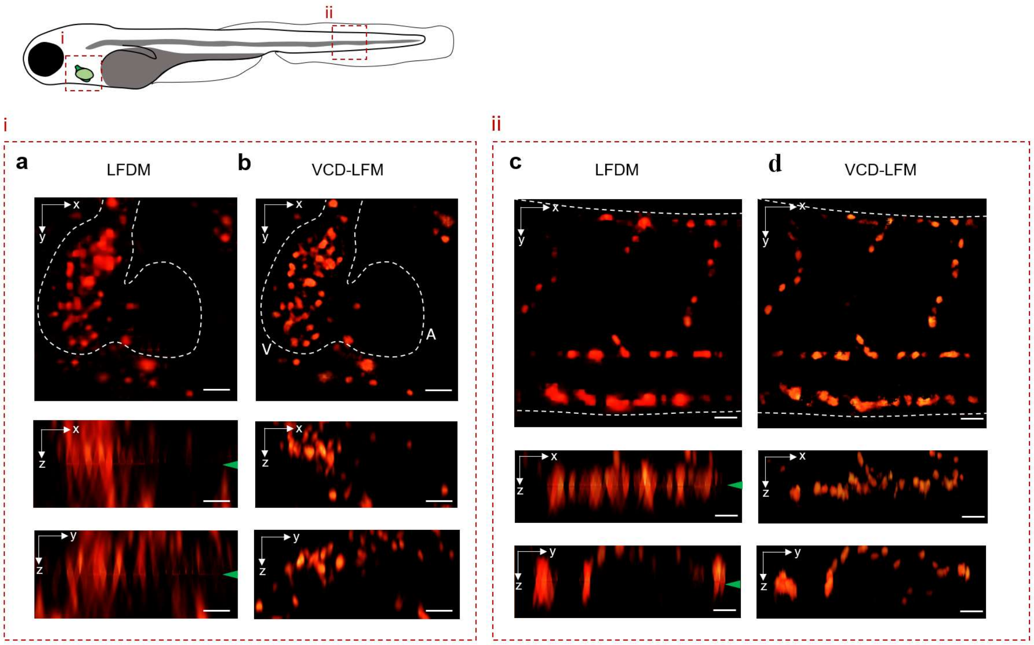


**Supplementary Figure 14**

**Reconstruction of red blood cells in the zebrafish embryo circulation system (heart and tail, *Tg(gata1:dsRed)*, 4dpf).**

(**a**),(**b**) MIPs of the reconstructed RBCs in the ventricle region, by LFDM and our VCD-LFM, respectively. VCD-LFM eliminates the artefacts at the focal plane (green arrows) and shows improved spatial resolution. Dotted line denotes the anatomy of the zebrafish heart. A, atrium; V, ventricle. (**c**),(**d**) Reconstructed RBCs in the tail vessels. Scale bar, 30 μm.

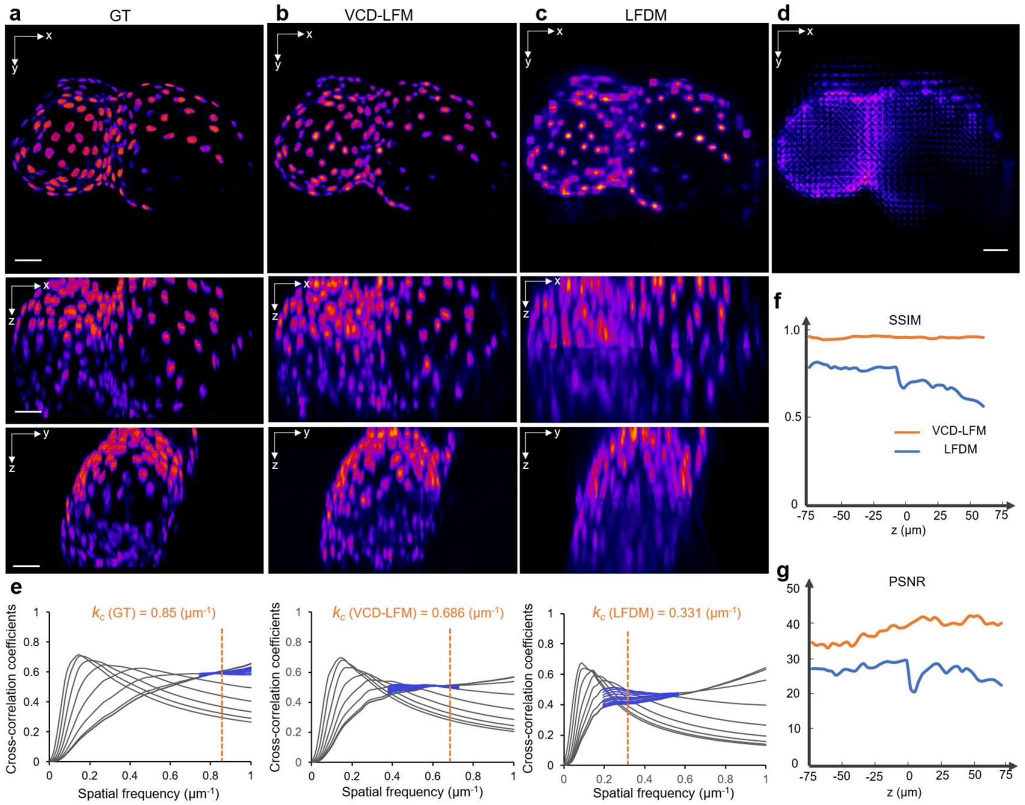


**Supplementary Figure 15**

**Comparison between confocal ground truth, VCD-Net reconstruction and LFD reconstruction of cardiomyocyte nuclei.**

(**a**) MIPs of the ground truth image (acquired by confocal microscope) of anaesthetized nucleus-labeled heart (*Tg(myl7:NLS-EGFP),* 4dpf). (**b**),(**c**) Reconstructions of the synthetic light-field image (**d,** from LFP of **a**) by our VCD-LFM and LFDM, respectively. It's visually obvious that VCD-LFM can recover the structures more accurately than those from LFDM, as compared to the ground truth. Furthermore, to quantify the achieved resolution and fidelity of VCD-Net and LFD, we calculated the cut-off frequency $k_c$ , the SSIM and PSNR indices of the reconstructed images. (**e**) We estimated the decorrelation functions and calculated the cut-off frequency $k_c$ values for the GT, VCD-LFM and LFDM images (xz MIPs in **a**-**c**), which here corresponded to the resolutions of 1.6, 1.983, and 4.1 μm in GT, VCD-LFM and LFDM images, respectively. (**f**),(**g**) The SSIM and PSNR indices across the depth of VCD-Net and LFD results, respectively. Scale bar, 30 μm.

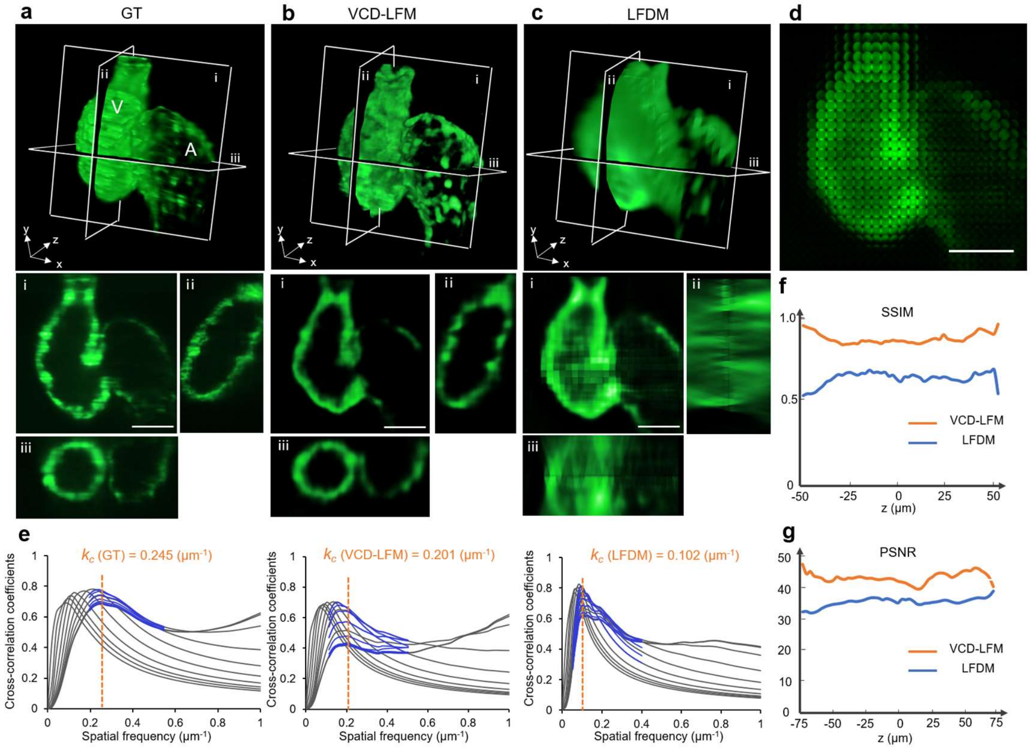


**Supplementary Figure 16**

**Comparison between HR light-sheet ground truth, VCD-Net reconstruction and LFD reconstruction of densely labelled trabecular myocardium.**

(**a**) 3D rendering of high-resolution image (acquired by light sheet fluorescence microscopy with retrospective gating) of myocardium (*Tg(cmlc2:gfp),* 4dpf). A: atrium; V: ventricle. (**b**),(**c**) 3D light-field reconstructions by VCD-LFM and LFDM, respectively. i to iii show the planes of ventricle and atrium indicated in the 3D renderings. As compared to the LFDM result showing obvious artefacts and ambiguous resolution, VCD-LFM successfully reconstruct the continuous myocardium with accurate shape recovered. (**d**) The light-field projection image used for reconstruction. (**e**) The decorrelation functions and the corresponding cut-off frequency $k_c$ quantitatively indicated the resolutions of 5.6, 6.7, and 13.3 μm in GT, VCD-LFM, and LFDM images (yz planes in **a**-**c**). (**f**),(**g**) The SSIM and PSNR indices across the depth of the reconstructed VCD-LFM and LFDM images, with referring to the light-sheet ground truth. Scale bar, 50 μm.

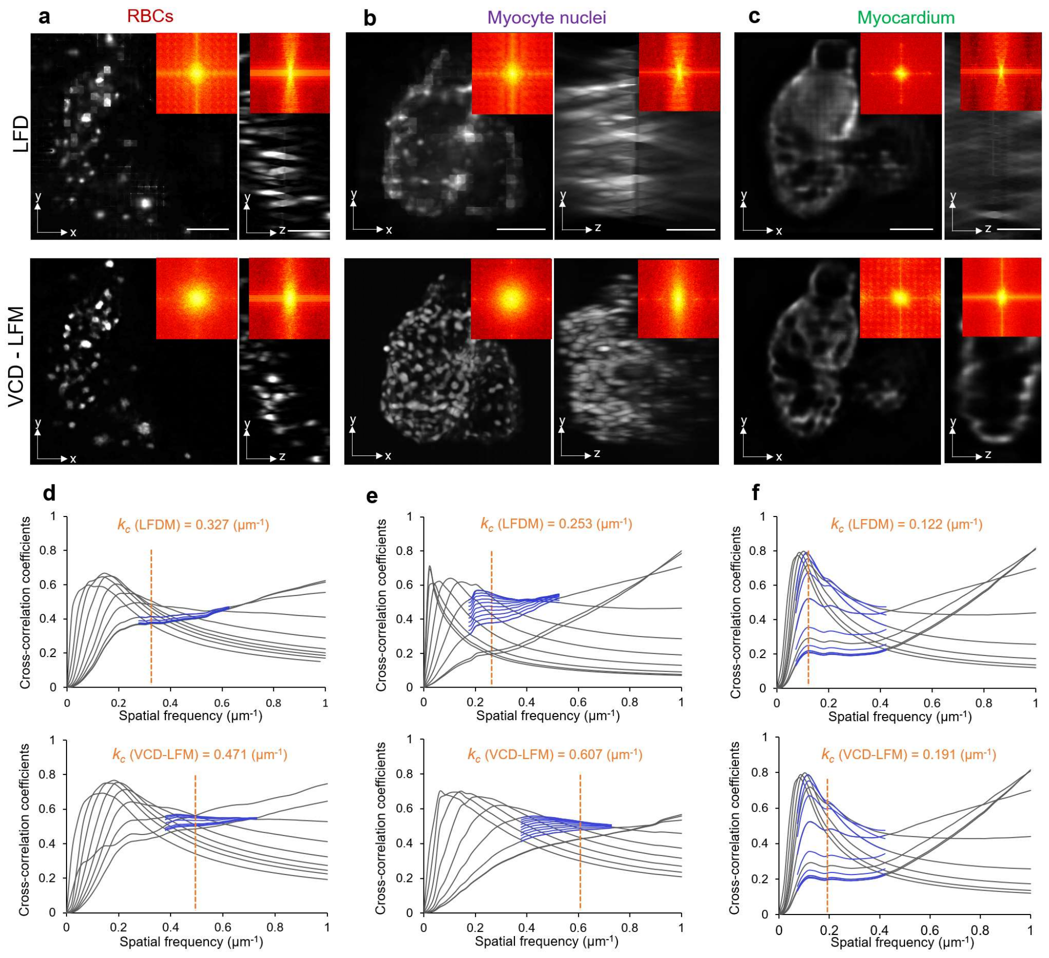


**Supplementary Figure 17**

**Resolution quantification based on experimental cardiac data.**

(**a**), (**b**) MIPs of RBCs and myocyte nuclei reconstructed by LFD (upper row) and VCD-Net (lower row). (**c**) Planes of dense myocardium reconstructed by LFD (upper row) and VCD-Net (lower row). The insets in (**a**)-(**c**) show the corresponding Fourier spectrograms of GT, VCD-Net reconstruction and LFD reconstruction. (**d**), (**e**), (**f**) The decorrelation analysis of reconstructed images in yz planes of (**a**), (**b**), (**c**), respectively. gray lines, all high-pass filtered decorrelation functions; blue lines, decorrelation functions with refined mask radius and high-pass filtering range; dashed vertical line, the cut-off frequency $k_c$ that represents the derivative of the smallest resolvable distance. The calculated resolutions of LFDM in (**d**), (**e**), (**f**) are 4.2, 5.3, and 11.1 μm, respectively. These values are compared to resolutions of 2.8, 2.2, and 7.1 μm in (**d**), (**e**), (**f**) by VCD-Net. Scale bar, 50 μm.

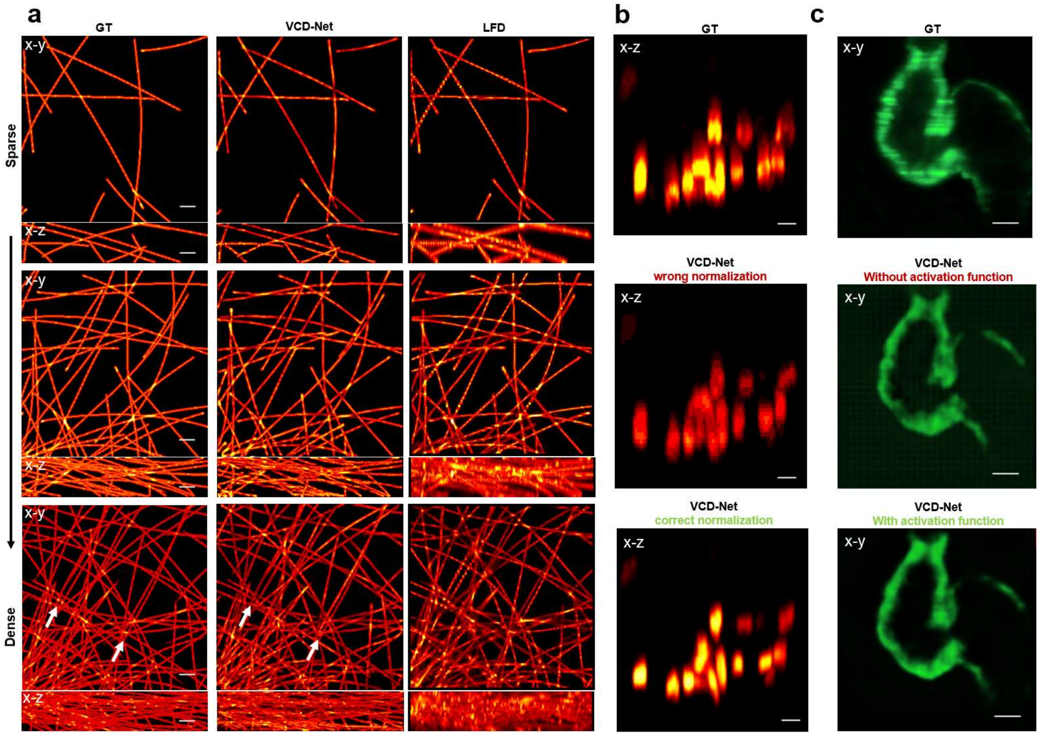


**Supplementary Figure 18**

**Failure cases of VCD-Net reconstructions.**

(**a**) Reconstructions of light-field inputs (synthetic tubulins) with different signal density. While VCD obviously showed better reconstruction quality than LFD at all three densities. Some artefacts containing inaccurate distribution or intensity (e.g., white arrows) arose when these line-like signals become highly dense. Scale bar, 50 μm (**b**) Abnormal dynamic range of recovered RBC signals, owing to inappropriate normalization applied in the network model. Given the various intensity of signals, normalizing images with fixed maximum value was not reliable enough, and occasionally caused poor reconstruction (middle). Normalizing the training data with dynamic maximum value (110% of the peak pixel intensity in input data) successfully circumvented this problem (bottom). Scale bar, 10 μm.(c) Grid-like fix pattern noises shown in the reconstruction by VCD-Net without a nonlinear activation function included (middle). A tanh activation function was added to the last layer of the upsampling path to regularize the output into the same intensity range of input ([-1 1]). This permitted a more efficient back-propagation to iteratively update the weights and biases of output layer, thereby better pushing the training outputs toward the ground truths. The tuned network substantially removed those hallucinations from the output (bottom). Scale bar, 50 μm.

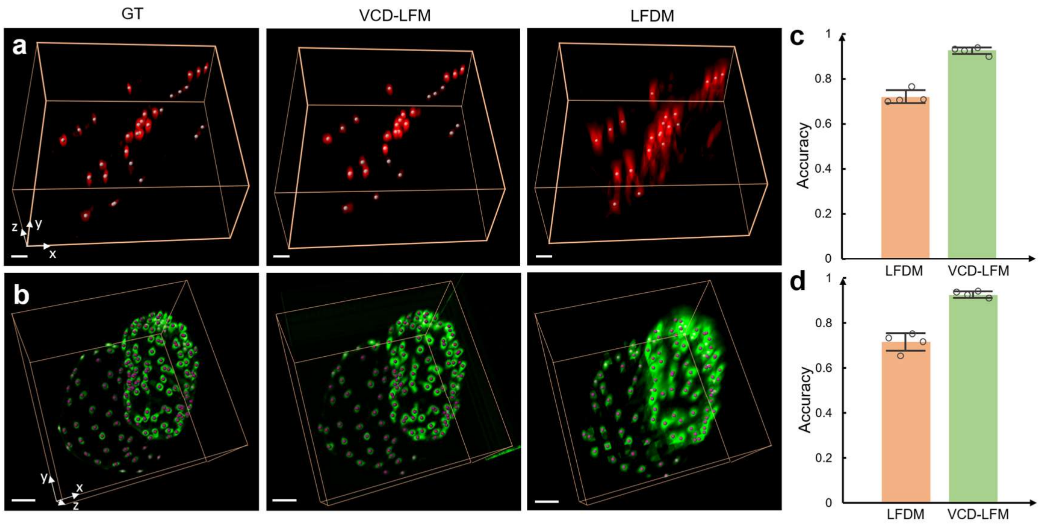


**Supplementary Figure 19**

**Accuracy of VCD-Net for resolving and counting individual RBCs and cardiomyocyte nuclei.**

(**a**) RBCs and (**b**) Cardiomyocyte nuclei three-dimensionally visualized using confocal (GT), VCD-LFM and LFDM results, as shown in left, middle and right column, respectively. The individual RBCs/cardiomyocyte nuclei resolved by three methods were further segmented and counted using Imaris (gray and purple dots). (**c**),(**d**) The single-cell counting accuracy, which is defined as the (VCD-LFM number / GT number) × 100% or (LFDM number / GT number) × 100%. As compared to the standard counting results based on HR ground truth, the averaged counting accuracy based on LFDM is at an average of 0.72 ± 0.028 in red blood cells and 0.72 ± 0.04 in cardiomyocytes nuclei, while this value is increased to 0.92 ± 0.014 and 0.92 ± 0.014 for VCD-LFM. Error bars indicate standard deviation. Experiments were repeated for 4 groups of zebrafish data. Scale bar, 20 μm.

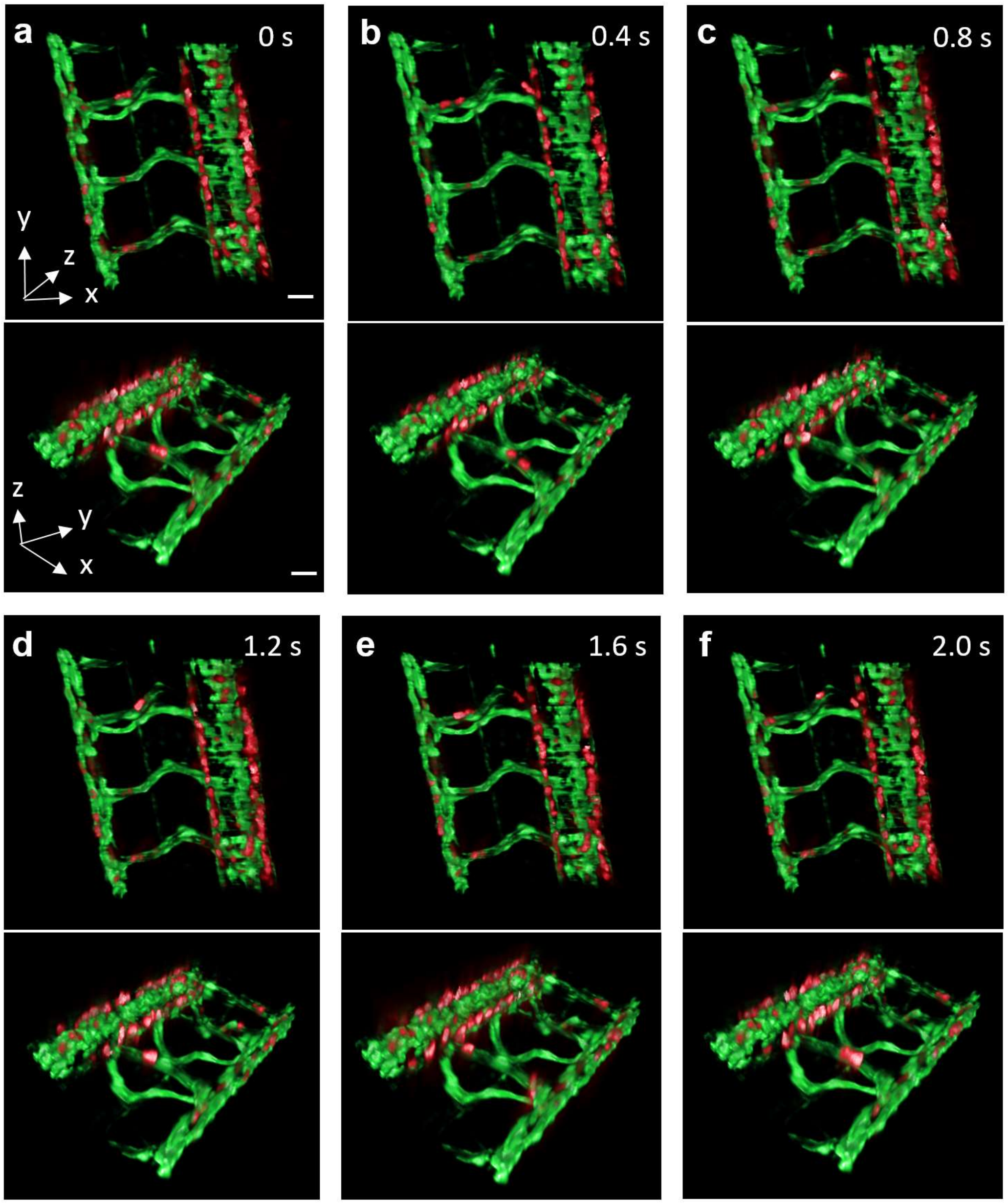


**Supplementary Figure 20**

**5D (3D space + time + spectrum) imaging of the blood flow in the tail veins of zebrafish embryo by the combination of VCD-LFM with SPIM.**

The red LFM channel visualizes the flowing RBCs while green SPIM channel visualizes the static vein. (**a**)-(**f**) The time sequence revealing how the RBCs flowing through the intersegmental veins. Scale bar, 30 μm. More visualization details and the quantitative analysis of the 3D blood velocity distribution in the vein system can be found in **Supplementary Video 7, 8**.

#### Supplementary Note 1: Model of light-field projection

The light-field projection can be modeled through scalar diffraction theory[4,5]. A point source in object space is projected onto the imaging sensor, with a complex 2D diffraction pattern encoding the 3-D position information of the point source. A library of Point Spread Functions (PSFs) can be calculated to characterize this projection process. Compared to conventional microscope, the light-field PSFs map the transformation from 3D object into a 2D plane and are also spatially variant, thereby a unique PSF is considered for each point in the region of interest.

A linear forward model can be applied to simulate the formation of light-field image

$$\boldsymbol{f}(\boldsymbol{x}) = \int |\boldsymbol{h}(\boldsymbol{x},\boldsymbol{p})|^2\, \boldsymbol{g}(\boldsymbol{p})\, \boldsymbol{dp} \tag{1}$$

where $\boldsymbol{p} = (\boldsymbol{p_1}, \boldsymbol{p_2}, \boldsymbol{p_3})$ denotes the position of the point source in object space whose intensity is defined as $g(\boldsymbol{p})$, and $\boldsymbol{x} = (\boldsymbol{x_1}, \boldsymbol{x_2})$ represents the coordinates on sensor plane. The optical impulse response $h(\boldsymbol{x}, \boldsymbol{p})$ is a transfer function derived from the wave optic model of light-field microscope, where a squared modulus is used owing to the incoherence of fluorescence imaging. A 2D light-field raw image $f(\boldsymbol{x})$ is finally calculated from 3D object $\boldsymbol{p}$.

Then, a discrete model can be used in implementation

$$\boldsymbol{f} = H \cdot \boldsymbol{g} \tag{2}$$

Where vector $\boldsymbol{f}$ represents the light-field raw image we will generate, and vector $\boldsymbol{g}$ is the discrete volume in form of high-resolution (HR) 3D image stack. The coefficients $H$ is the PSF library we generated using code[5].

The accuracy of this light-field projection is crucial for the VCD-Net to implement reliable data training, and accurate inference afterwards. Therefore, we evaluated the similarity between the synthetic (projection from HR data) and experimental (acquired using LFM) light-field raw images of the same sub-diffraction beads. In **Supplementary Note Fig. 1**, we showed the synthetic and measured PSFs of the identical beads located at different depths (-15 µm to 15 µm), and quantitatively compared them via calculating the PSNR (Peak Signal-to-Noise Ratio) and SSIM (Structural Similarity). The synthetic light fields by light-field projection were thus verified to be enough accurate for VCD-Net training.

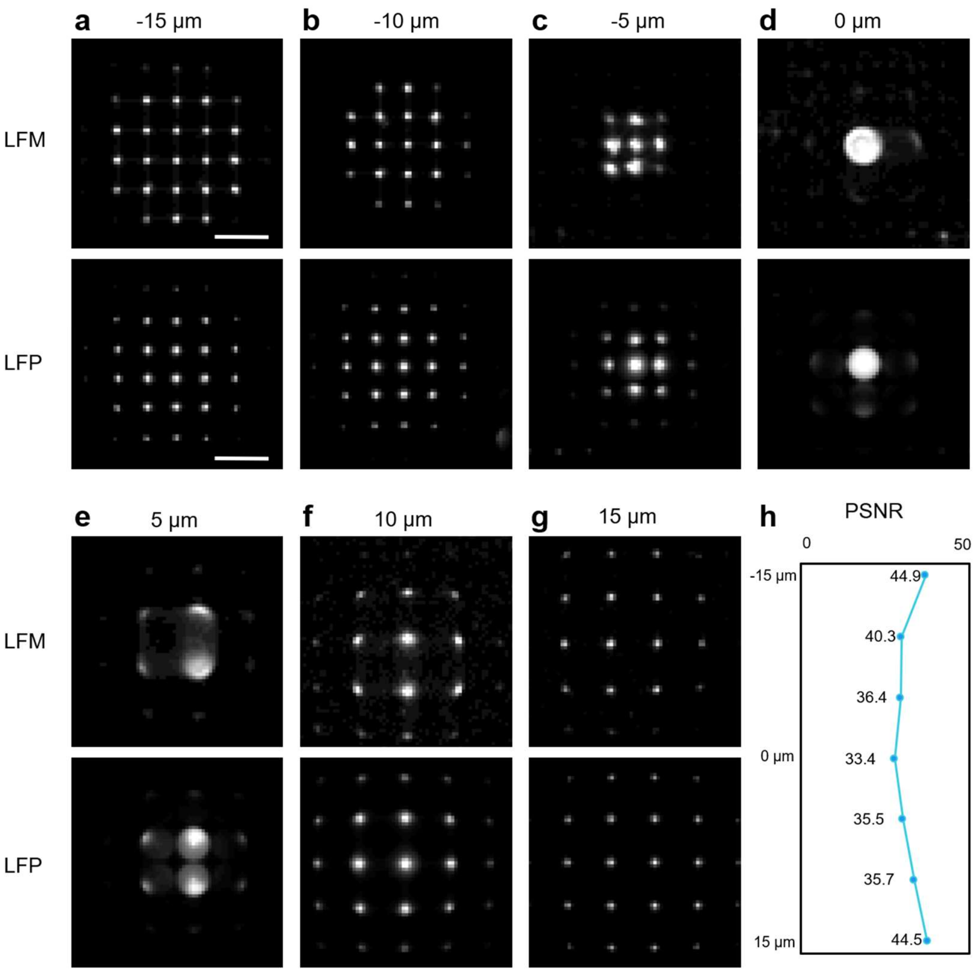


**Supplementary Note Figure 1. Comparison between synthetic and experimental light-field raw images.** (**a**)-(**g**) The light-field raw images of identical sub-diffraction beads (z = -15 μm to 15 μm) by experimental LFM (up) and synthetic LFP (light-field projection) (bottom), respectively. (**h**) The PSNR indices of the synthetic and experimental light-field images. The high-level values (greater than 30) indicate the high similarity between the synthetic and experimental data, quantitatively proving the accuracy of the light-field projection model. Scale bar, 5 μm.

#### Supplementary Note 2: VCD-Net implementation details

The VCD-Net is designed to reconstruct 3D volumes directly from 2D light-field raw images. It includes 2 stages: 1. VCD training based on high-resolution 3D images and their synthetic light-fields; 2. Inference on experimental light-field images. Training is required only once for all the following inferences, or it can be skipped if a well-trained VCD-Net has been previously established.

To prepare training data, we used light-field projection (**Supplementary Note 1**) to convert the high-resolution 3D image stacks into corresponding synthetic light-fields. Required by the light-field projection, the light-field PSFs $\boldsymbol{H}$ should be defined at the beginning, based on both the system setup (the magnification and NA of the objective, the pitch and focal length of the MLA, etc.) and the experiment specifications (the pixel size of the synthetic light-fields, the reconstruction range and z step size, etc.). PSFs were computed using *computePSF_GUI.m* in code[5]. We then resampled and cropped them in compliance with the voxel size and reconstruction range that are specified by PSFs $\boldsymbol{H}$. They were next augmented via random rotation, shift and flip to increase the data scale to avoid overfitting. Through light-field projection, these high-resolution volumes were corresponded to their synthetic light fields. We cropped these image pairs into small patches of identical size (e.g. 176 × 176 × 31 px and 176 × 176 px) and blank patches with scant information were discarded using a criterion of the statistic mean and variance of the pixel intensities.

In the training stage, the VCD-Net iteratively minimized the difference between the intermediate outputs and high-resolution 3D image stacks. We chose an Adam optimizer with decaying learning rate and Mean Square Error as loss function. For every 10 iterations, we saved the reconstruction results of validation dataset (a small portion of data which we manually selected and excluded from the training at the beginning). The performance was evaluated on both the loss function and the visual examination of the validation results. The model with best performance was chosen for experimental data inference.

Before inference, the experimental light-field raw images were resampled according to the same PSF configurations in training stage. The resampling was done by *ImageRectification_GUI.m* in code[5]. The size of input light-field raw image doesn't necessarily have to be the same as training patches. It's also recommended to adjust the input images to have dynamic range similar with the training patches. The inference step can be very quick, and the output will have the same voxel size and depth range as PSFs $\boldsymbol{H}$ specified.

The detailed VCD-Net model implementation is listed in **Supplementary Table 1**. The computation environment and cost are given in **Method**. Regarding the network performance, it should be noted that with using a data-driven model, the reconstruction quality of our VCD-Net heavily relies on the training data. In our experiments, though confocal microscopy has generated high-resolution 3D images (*C. elegans*, zebrafish) that have permitted successful VCD-Net training and inference, their results were still not ideal owing to the inevitable aberration and scattering from real samples. We also used synthetic dataset (fluorescence beads, tubulins), for which we can control the quality via blurring and noise models. We trained the VCD-Net with this synthetic data and obtained reconstruction of various quality. The results shown in **Supplementary Fig. 7** implied that the network performance could be possibly optimized by using higher-quality training data. This indicates that VCD-Net reconstruction of biological specimens with quality better than those we have shown in this paper could be potentially realized through training the network with higher-quality data obtained by more ideal imaging techniques.

#### Supplementary Note 3: Comparison between VCD-Net and LFD + deep-learning image restoration

We further compared our 2D-3D VCD-Net reconstruction with a 2D-3D-better3D procedure accomplished by LFD reconstruction combined with state-of-the-art deep-learning image restorations (iso-CARE, 3D-CARE)[6]. The computation workflows, including the data training and image application, of VCD-Net, LFD + iso-CARE, and LFD + 3D-CARE were illustrated in below **Supplementary Note Fig. 3a.** The sparse point-like cardiomyocyte nuclei and densely-labeled continuous myocardium reconstructed by these three procedures were visually and quantitatively compared in **Fig. 3b**, **c** and **d**, **e**, respectively**.** Though LFD + iso-CARE didn't necessarily require HR labelling data for training, this strategy showed limited enhancement to LFD, for either myocyte nuclei or myocardium (**3b3, 3d3**), owing to the still suboptimal lateral quality of LFD result (**3b2, 3d2**). Unsurprisingly, the addition of 3D-CARE, which was trained by HR labelling data and corresponding synthetic LFD images, showed more significant improvement to LFD results (**3b4**, **3d4**). However, while LFD + 3D-CARE could reconstruct non-dense myocyte nuclei signals with similar quality (**3b4**, **3b5**) except for a few lost signals near the focal plane (white boxes), its recovery on densely-labeled myocardium was still far worse, as compared to VCD-Net (**3d4, 3d5**). Noticeable restoration hallucinations arose from the excessive artefacts in LFD results, and also caused unacceptably low structure similarity (SSIM) to the ground-truth data (**3e**).

It's also noted that while these combinatorial recovery strategies show more or less enhancement to LFD results, they have issue of low application efficiency. Since they need to first go through the iterative LFD for each frame of a light-field video, the processing speed is even lower than LFD only, over three-order-lower than VCD reconstruction, and thus impractical for many applications (**Supplementary Note Fig. 3f**). Furthermore, such additional deep-learning restoration to LFD results also requires a lot of LFD data prepared for model training, which is more time consuming than generating LFP data for VCD-Net training (~10× slower, **Supplementary Note Fig. 3f**). Therefore, from the perspectives of reconstruction quality, reconstruction speed and model implementation, which are all important factors for a computational imaging technique, we have shown the significant advantages of our VCD-Net approach, even when compared with the combination of two established restoration methods. Especially under dense labelling conditions, only VCD-Net can realize accurate light-field reconstruction at high resolution, which remains unachievable by alternative approaches.

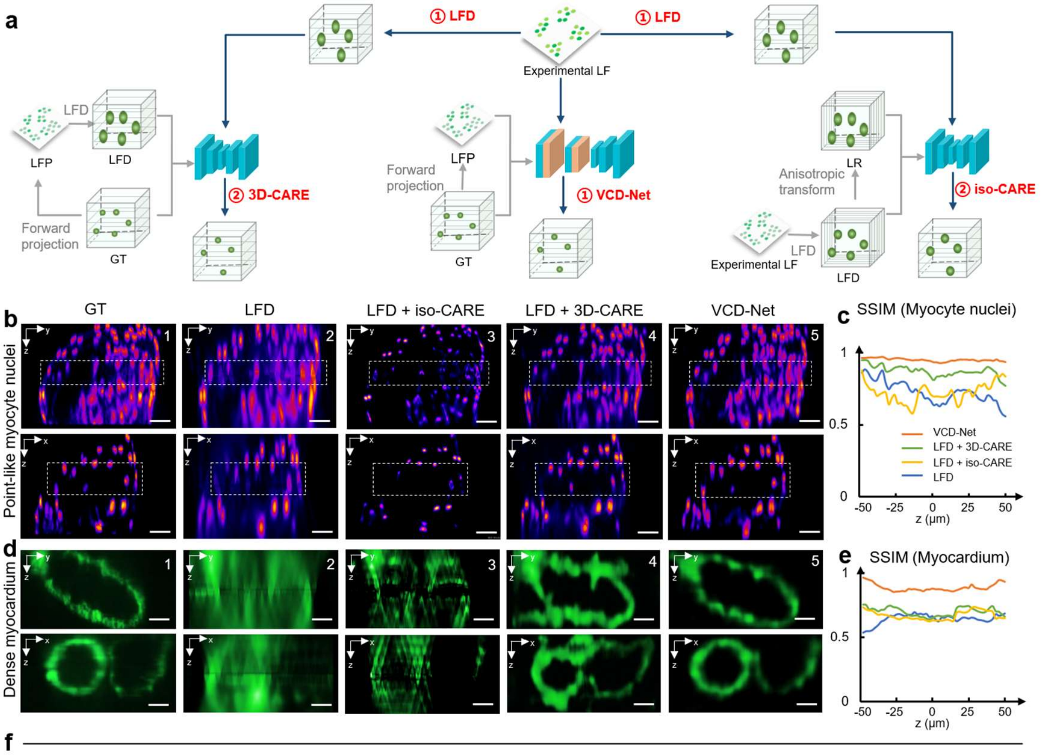


f

| Methods | Data preparation time | Model training time | Reconstruction speed |
|---|---|---|---|
| LFD | / | / | ~90*s* per volume |
| LFD + iso-CARE | ~20*h* (LFD) | ~3.3*h* | ~90.4*s* per volume |
| LFD + 3D-CARE | ~22.2*h* (LFD + LFP) | ~9.4*h* | ~90.5*s* per volume |
| VCD-Net | ~2.2*h* (LFP) | ~5*h* | ~0.03*s* per volume |

**Supplementary Note Figure 3. Comparison between VCD-Net and LFD + deep-learning image restoration.** (**a**) Workflows of LFD + 3D-CARE (left), VCD-Net (middle) and LFD + iso-CARE (right) reconstruction procedures. (**b**) Comparative MIPs in *y-z* (top) and *x-z* (bottom) planes of the labeled cardiomyocyte nuclei by confocal microscope (GT, b1), LFD (b2), LFD plus iso-CARE (b3), LFD plus 3D-CARE (b4), and our VCD-Net (b5) from left to right, respectively. White boxes show the regions near the native focal plane. (**c**) Structure similarity (SSIM) curve across the depth of the reconstructions, indicating the accuracy of each method when reconstructing non-dense nuclei signals. (**d**) Comparative MIPs in *y-z* (top) and *x-z* (bottom) planes of the densely-labeled myocardium by light-sheet microscope (GT, d1), LFD (d2), LFD plus iso-CARE (d3), LFD plus 3D-CARE (d4), and our VCD-Net (d5) from left to right, respectively. (**e**) SSIM curve of each method when reconstructing highly-dense myocardium signals. Scale bar, 25 μm. (**f**) Table comparing the efficiency the 4 light-field reconstruction strategies, in term of the model implementation time and reconstruction speed. The data preparation and model training time of each method was calculated based on the same 800 groups of myocyte data. The reconstruction time of each approach was calculated by recovering a 3D output of myocytes (330*330*51) from the same light-filed input (330*330). All the tests were performed on the same workstation equipped with RTX 2080 Ti graphic card.

#### Supplementary Note 4: The generalization ability of VCD-Net

Generalization ability has been an important characteristic of a deep-learning-based model. In this section, we demonstrate hybrid data training/multi-sample recovery, and cross-sample/transfer learning applications, to evaluate the generalization ability of VCD-Net.

##### 1. Performance of VCD-Net trained on hybrid cardiac data

Unlike the conventional network implementation based on a single type of sample data, here a hybrid cardiac VCD-Net was first trained on mixed datasets containing GT-LFP image pairs of myocyte nuclei, red blood cells and myocardium (**Supplementary Note Fig. 4.1a**), and then applied to the light-field reconstruction of all the three types of samples. The results by such a hybrid VCD-Net (**4.1c**) were compared with those by three individual VCD-Nets that were trained on myocytes nuclei, RBCs and myocardium separately (**4.1b**).The similar reconstruction quality by the hybrid network indicates that the VCD-Net could generalize well when trained on many datasets and applied to many samples.

##### 2. Cross-sample and transfer learning applications of VCD-Net

We trained 3 VCD-Nets, using image pairs of the sparse red blood cells, mid-density myocyte nuclei, and highly-dense myocardium, respectively, and applied each of networks to the reconstruction of all the three types of samples (**Supplementary Note Fig. 4.2).** As compared to the high-quality reconstruction for the same types of samples (**a1**, **b1**, **c1**), the quality of cross-applications was also acceptable (**a2**, **b2**) when recovering the similar types of signals (point-like myocyte nuclei network for point-like RBCs in **a2**, or *vice versa* **b2**). At the same time, the reconstruction quality was severely compromised when applying point-signals-trained networks to continuous myocardium data (**a3 middle**, **b3 middle**), or *vice versa* (**c2 middle**, **c3 middle**).

To overcome the barrier between such different signal types, we also introduced transfer learning, which leveraged the knowledge already learned by the previously-trained network and thus required much fewer training data and iterations, to further enhance the generalization ability of VCD-Net. In practice, we saved the best checkpoints of the pre-trained point-signal-based and continuous-signal-based VCD models, and then trained them using small amount (~20%) of data from continuous sample and point-like samples, respectively. As shown in the right columns of **a3**, **b3**, **c2**, **c3**, after transfer learning applied, the previously corrupted reconstructions caused by style-mismatching were significantly mitigated. Sufficiently accurate reconstructions have been provided, as compared to the GT data. Therefore, the VCD-Net could be highly generalizable when trained on one type of data and applied to another, especially when a transfer learning based on small amount of target data involved.

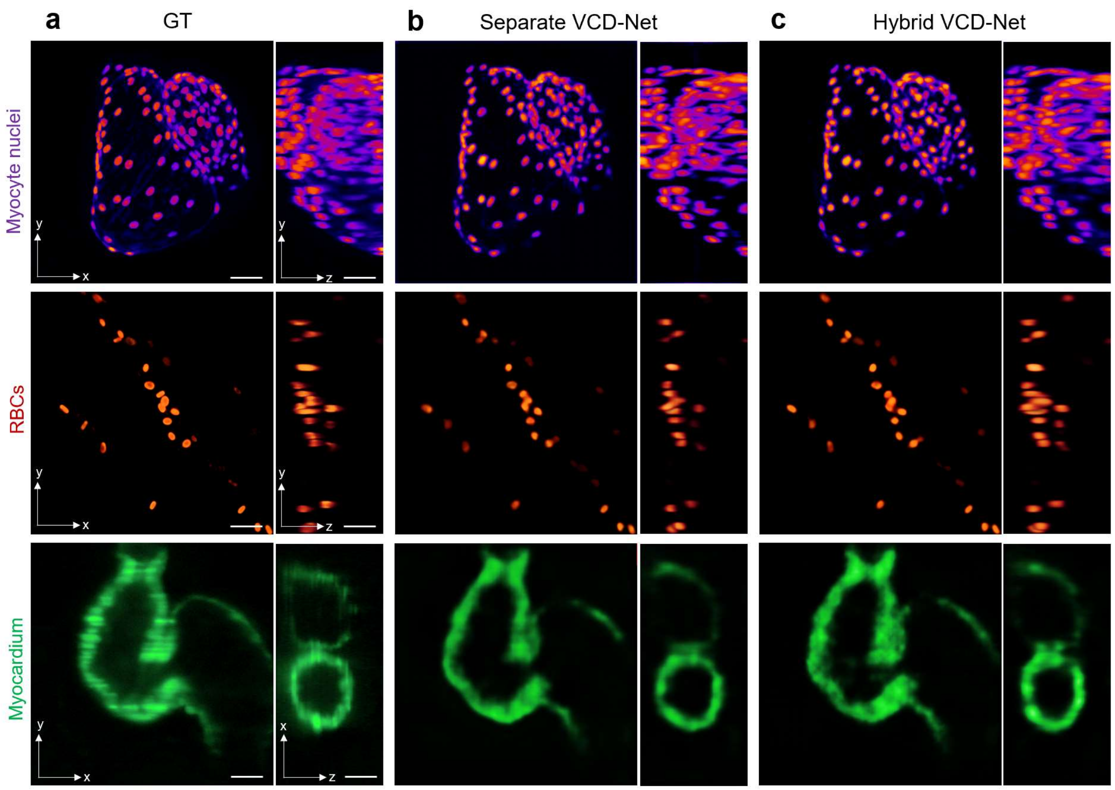


**Supplementary Note Figure 4.1. Performance of VCD-Net trained on hybrid cardiac data.** (**a**) High-resolution confocal images of the myocyte nuclei (row 1), RBCs (row 2), and light-sheet microscopy image of myocardium (row 3). (**b**),(**c**) Reconstructions of the same samples by separate VCD-Nets with each trained on a single type of data, and hybrid VCD-Net trained on mixed cardiac datasets, respectively. It's shown that the myocyte nuclei, RBCs and myocardium all can be reconstructed using a single hybrid VCD-Net. Scale bar, 30 μm.

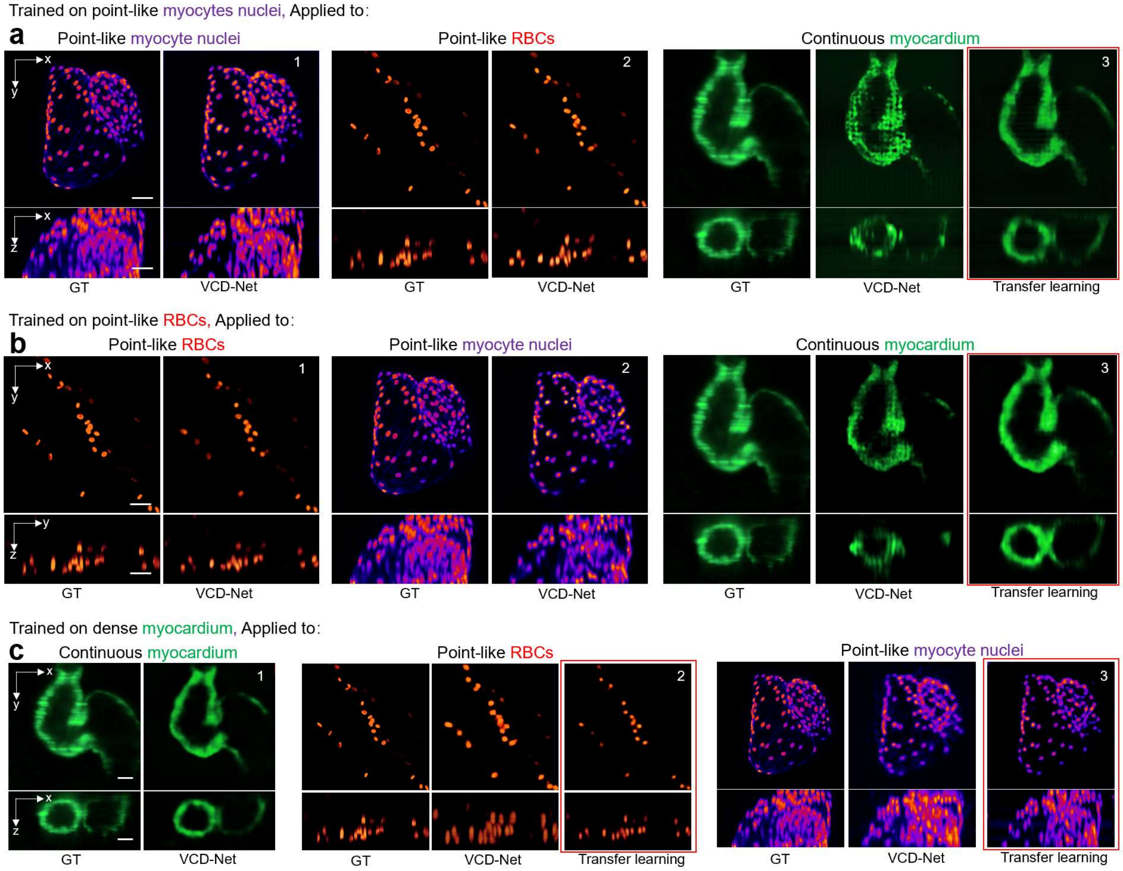


**Supplementary Note Figure 4.2. Cross-sample and transfer learning applications of VCD-Net. (a)-(c)** Cross applications to myocyte nuclei, RBCs, and myocardium, by VCD-Net models which are solely trained on myocyte nuclei, RBCs, and myocardium, respectively. It's shown that while two models trained on point-like myocyte nuclei and RBCs can generalize well to each other, they both show poor results on the dense myocardium, and *vice versa*. This limitation from very different signal types can be further overcome by introducing transfer learning (red boxes). Scale bar, 30 μm.

#### Supplementary Note 5: Pipeline for behavior analysis of *C. elegans* (motor neuron)

Unlike previous works that require extra bright-field images to analyze acting worm's locomotion[7], we developed a 2D fluorescence image-based pipeline to readily analyze the speed and posture of the worm (**Supplementary Note Fig. 5a**). The fluorescence images were first reconstructed by VCD-Net and 2D MIPs were used for the following analysis. Each 2D projection was then enhanced by histogram equalization to highlight weak worm outline. Worm body was segmented, and the edge was computed using Canny filter. We calculated the Euclidean distance transform inside the worm body on the edge image and the ridge of distance transformation thus indicated the center line of the animal. Due to the weak signals, there's variance shown in segmentation results at worm head and tail. To avoid inconsistence of the start points in the extracted center lines, we tracked two neurons near the head and adopted their positions as the seed in searching algorithm: we initialized our searching kernel around seed neurons and seek for points that lie on the center line with identical spacing from head to tail. Once we obtained the center line $P(i,t) = \big(x(i,t)\,,\ y(i,t)\big)$ of the worm at different time, we followed the previously-established mathematical definitions[4] to quantify the curvatures and moving speed. Here, the $P(i,t)$ denotes a point at time $t$ with index $i$ among all the points $(1,2 \ldots n)$ at the center line. Each point $P(i,t)$ is associated with the radius of the body at that location and the curvature $\kappa(i,t)$ can be thereby calculated by following formula

$$\kappa(i,t) = \frac{x'(i,t)y''(i,t) - x''(i,t)y'(i,t)}{\big(x'^2(i,t) + y'^2(i,t)\big)^{\frac{3}{2}}}$$

Meanwhile, the travel speed can be expressed as

$$mean\{\frac{\partial}{\partial \mathrm{t}}P(i,t), i = 1,2 \ldots n\}$$

We adopted a 2D U-Net for high-throughput, automatic worm segmentation (**Supplementary Note Fig. 5b**). We manually annotated 114 images (MIPs). The segmentation model was trained between the histogram equalized images and our annotations. The input images were resized to 256 x 256 pixels and augmented via rotation, shifting, flipping and zooming to avoid overfitting. We used Adam optimizer and binary cross-entropy as loss function. The model was built with Keras and trained on a Nvidia GeForce RTX 2080 Ti graphic card. The training typically takes ~15 minutes.

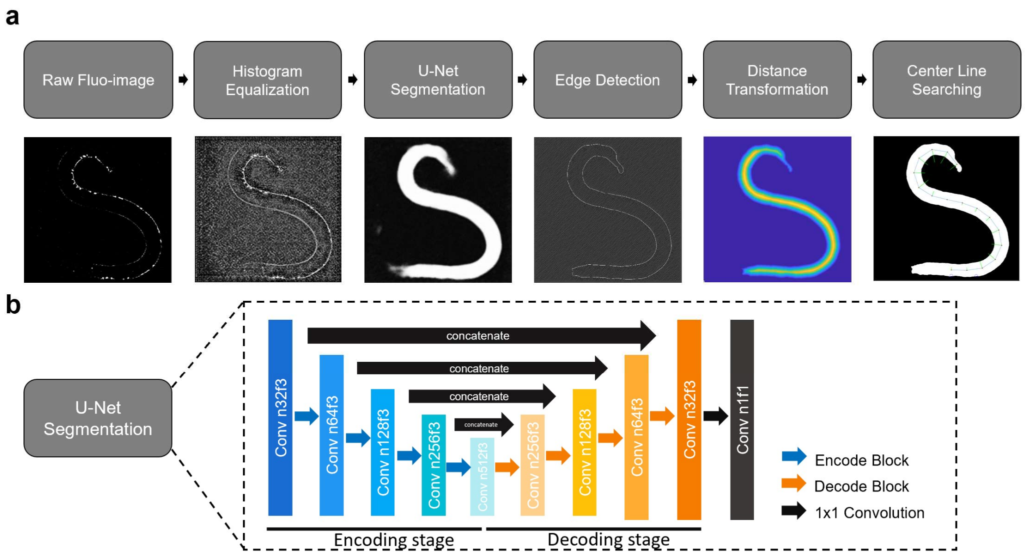


**Supplementary Note Figure 5. Pipeline for behavior analysis of C. elegans.** (**a**) Steps of image processing to extract center line from raw fluorescence image for behavior analysis. The body curvature and movement velocity were then inferred from the center line. (**b**) Structure of segmentation U-Net. Color blocks represent 2D convolution/transposed convolution layers with parameters n (channel number) and f (filter size). Each encoder block contains 2 convolution layers and each decoder block contains 1 transposed convolution layer followed by 2 convolution layers. The final black block has only one convolution layer. We used 2D max pooling for image down-sampling and transposed convolution with stride 2 for image up-sampling.

### Supplementary Table 1. Detailed Architecture of the VCD-Net

| Alias | VCD-Net - structural | VCD-Net - functional |
|---|---|---|
| | Conv n128f7s1 | Conv n128f5s1 |
| | Subpixel conv | Subpixel conv |
| | Conv n64f3s1 | Conv n64f3s1 |
| | Subpixel conv | Subpixel conv |
| | Conv n32f3s1 | Conv n32f3s1 |
| | Subpixel conv | Subpixel conv |
| | Conv n16f3s1 | Conv n16f3s1 |
| | Subpixel conv | Subpixel conv |
| | Conv n8f3s1 | Conv n8f3s1 |
| | Conv n8f3s1 | |
| | Batch normalization | |
| | ReLU | |
| | Conv n64f3s1 | Conv n64f3s2 |
| | Batch normalization | Leaky-ReLU |
| l1 | ReLU | Conv n64f3s2 |
| | Conv n128f3s1 | |
| | Batch normalization | |
| l2 | ReLU | |
| | Add [l2, l1] | Leaky-ReLU |
| l3 | Max pooling f3s2 | Conv n128f3s2 |
| | Conv n256f3s1 | |
| | Batch normalization | |
| l4 | ReLU | |
| | Add [l4, l3] | Leaky-ReLU |
| l5 | Max pooling f3s2 | Conv n256f3s2 |
| | Conv n512f3s1 | |
| | Batch normalization | |
| l6 | ReLU | |
| | Add [l6, l5] | Leaky-ReLU |
| l7 | Max pooling f3s2 | Conv n512f3s2 |
| | Conv n512f3s1 | |
| | Batch normalization | |
| l8 | ReLU | |
| | Add [l8, l7] | |
| l9 | Max pooling f3s2 | |
| | Conv n512f3s1 | |
| | Batch normalization | |
| l10 | ReLU | |
| | Add [l10, l9] | |
| | Max pooling f3s2 | |
| l11 | Up sampling | ReLU |
| | | Up sampling |
| | | Conv n512f3s1 |
| | Concat [l9, l11] | Concat [l7, l11] |
| | Conv n512f3s1 | |
| | ReLU | ReLU |
| | Batch normalization | Up sampling |
| l12 | Up sampling | conv n256f3s1 |
| | Concat [l7, l12] | Concat [l5, l12] |
| | Conv n512f3s1 | |
| | ReLU | ReLU |
| | Batch normalization | Up sampling |
| l13 | Up sampling | conv n128f3s1 |
| | Concat [l5, l13] | Concat [l3, l13] |
| | Conv n256f3s1 | |
| | ReLU | ReLU |

| | | |
|---|---|---|
| | Batch normalization | Up sampling |
| l14 | Up sampling | Conv n64f3s1 |
| | Concat [l3, l14] | Concat [l1, l14] |
| | Conv n128f3s1 | |
| | ReLU | ReLU |
| | Batch normalization | Resize |
| l15 | Up sampling | Conv n31f3s1 |
| | Concat [l1, l15] | |
| | Conv n51f3s1 | |
| | ReLU | |
| | Batch normalization | |
| | Resize | Conv n31f3s1 |
| | Tanh | ReLU |

There are two variations for the architecture of the VCD-Net. The VCD-Net-structural handles structural signals (e.g., beads, red blood cells, myocyte nuclei and myocardium) and the VCD-Net-functional) deals with functional imaging (e.g., calcium signals of *C. elegans* neurons) using a smaller model size and higher inference speed. "Conv" is the abbreviation for convolutional layer, the parameters of which are number of output channels (n), filter size (f) and stride (s). "Subpixel conv" is abbreviation for subpixel convolutional layer that up-scales the size of the feature maps by a factor 2. The "Up sampling" layer also up-scales the size of the feature maps by a factor 2 using bilinear (in VCD-Net-structural) or nearest neighbor (in VCD-Net-functional) interpolation. The "Resize" layer resizes the feature maps to the same size with the light-field image. "Add" is the element-wise add function. "Concat" is abbreviation for concatenation layer, which combines the feature maps of the two input layers along the "channel" dimension.

### Supplementary Table 2. Experiments setup

| Experiment | Beads | Worm (motor neuron) | Worm (pan neuron) |
|---|---|---|---|
| Magnification / NA | 40× / 0.8 | 40× / 0.8 | 40× / 0.8 |
| Diffraction index | 1.33 (water) | 1.33 (water) | 1.33 (water) |
| MLA focal length | 3500 µm | 3500 µm | 3500 µm |
| MLA pitch / Nnum[1] | 150 µm / 11 | 150 µm / 11 | 150 µm / 11 |
| pixel (voxel) size | 0.34 × 0.34 (× 1) µm（Aniso net）or 0.34 × 0.34 (× 0.34) µm (Iso net) | 0.34 × 0.34 (× 1) µm | 0.34 × 0.34 (× 1) µm |
| z-range / z-step | - 30 µm to 30 µm / 1 µm (Aniso net) or 0.34 µm (Iso net) | - 30 µm to 0 µm / 1 µm | - 30 µm to 0 µm / 1 µm |
| OSR[2] | 3 | 3 | 3 |
| **Training stage** | | | |
| Training data | Synthesized 3-D beads | Static *C.elegans* (motor neuron) images from 20 samples (L4 stage) using confocal microscope | Static *C. elegans* (pan neuron) images from 24 samples (L4 stage) using confocal microscope |
| Training patch size | 176 × 176 and 176 × 176 × 61(Aniso net) or 176 x 176 × 177(Iso net) | 176 × 176 and 176 × 176 × 31 | 176 × 176 and 176 × 176 × 31 |
| VCD-Net model | VCD-Net-structural | VCD-Net-functional | VCD-Net-structural |
| **Inference stage** | | | |
| Sample | 0.5 µm fluorescent beads (excitation @ 540 nm, emission @ 580 nm) | ZM9128 *hpIs595[Pacr-2(s)::GCaMP6(f)::wCherry]* @ L4 Stage | QW1217 *hpIs467[Prab-3::NLS::RFP]* @ L4 Stage |
| Imaging setup | Supplementary Figure 1 | Supplementary Figure 1 | Supplementary Figure 1 |
| Original data size (row × col) | 2048 × 2048 (332.8 × 332.8 µm) | 2048 × 2048 (332.8 × 332.8 µm) | 2048 × 2048 (332.8 × 332.8 µm) |
| Exposure time | 10 ms @ 100fps | 2 ms @ 100fps | 20ms @ 50fps |
| Frames acquired | 1 | 6000 | 500 |
| Input data size (to VCD-Net) | 935 × 924 | 935 × 924 | 935 × 924 |
| Output data size | 935 × 924 × 61 (Aniso) or 177 (Iso) (317.90 × 314.16 × 60 µm) | 935 × 924 × 31 (317.9 × 314.16 × 30 µm) | 935 × 924 × 31 (317.9 × 314.16 × 30 µm) |
| Figures/Videos | Figure 1 c-g, Supplementary Figure 5-9 | Figure 2, Supplementary Figure 13, Video S2, S3 | Supplementary Figure 12, Video S1 |

| Experiment | Blood cells in zebrafish heart / tail | Myocardium in zebrafish heart | Myocyte nucleus in zebrafish heart |
|---|---|---|---|
| Magnification / NA | 20X / 0.5 | 20X / 0.5 | 20X / 0.5 |
| Diffraction index | 1.33 (water) | 1.33 (water) | 1.33 (water) |
| MLA focal length | 3500 µm | 3500 µm | 3500 µm |
| MLA pitch /Nnum | 150 µm / 11 | 150 µm / 11 | 150 µm / 11 |
| pixel (voxel) size | 0.68 × 0.68 (× 2) µm | 0.68 × 0.68 (× 2) µm | 0.68 × 0. 68 (× 3) µm |
| z-range / z-step | - 50 µm to 50 µm / 2 µm | - 50 µm to 50 µm / 2 µm | - 75 µm to 75 µm / 3 µm |
| OSR | 3 | 3 | 3 |
| **Training stage** | | | |
| Training data | Static RBC images in heart and tail from 16 fish (3dpf - 4dpf) using confocal microscope | Moving myocardium images in heart from 8 fish (4dpf) using light sheet microscope | Static nucleus images in heart from 23 fish (1dpf - 4dpf) using confocal microscope |
| Training patch size | 176 × 176 and 176 × 176 × 51 | 176 × 176 and 176 × 176 × 51 | 176 × 176 and 176 × 176 × 51 |
| VCD-Net model | VCD-Net-structural | VCD-Net-structural | VCD-Net-structural |
| **Inference stage** | | | |
| Sample | Zebrafish embryo *Tg(gata1:dsRed; cmlc2:GFP)* @ 3dpf / Zebrafish embryo *Tg*(gata1:dsRed; fi1:EGFP) @ 4dpf | Zebrafish embryo *Tg(cmlc2:gfp)* @ 4dpf | Zebrafish embryo *Tg(myl7:nls-EGFP)* @ 4dpf |
| Imaging setup | Supplementary Figure 2 | Supplementary Figure 2 | Supplementary Figure 2 |
| Original data size (row x col) | 768 x 768 (249.6 ×249.6 µm) / 1024 × 2048 (332.8 × 665.6 µm) | 768 × 768 (249.6 × 249.6 µm) | 768 × 768 (249.6 × 249.6 µm) |
| Original pixel size | 0.325 × 0.325 µm | 0.325 × 0.325 µm | 0.325 × 0.325 µm |
| Exposure time | 5 ms @ 200fps / 10 ms @ 100fps | 5 ms @ 200fps | 5 ms @ 200fps |
| Frames acquired | 450 / 600 | 450 | 450 |
| Input data size (to VCD-Net) | 341 × 341 / 462 × 946 | 330 × 330 | 330 × 330 |
| Output data size | 341 × 341 × 51(231.88 × 231.88 × 100 µm) / 462 × 946 × 51 (314.16 × 643.28 × 100 µm) | 330 × 330 × 51 (224.4 × 224.4 × 100µm) | 330 × 330 × 51 (224.4 × 224.4 × 100µm) |
| Figures/Videos | Figure 3 b-e, Supplementary Figure 14, 17a, 19a, 20, Video S4, S7, S8, S9 | Figure 3 h-k, Supplementary Figure 16, 17c, Video S6 | Figure 3 f, g, Supplementary Figure 15, 17b, 19b, Video S5 |

Nnum: the number of pixels behind each lenslet, used to resample/rectify the light-field raw images.

OSR: oversampling factor, used in light-field PSFs computation.

#### Supplementary Table 3. Reconstruction throughput of VCD-Net and LFD.

We ran a testbench using *C. elegans* dataset (**Supplementary Video 2**). Each group was reconstructed by both VCD-Net and LFD program using GPU and CPU, to compare the reconstruction throughput by two methods. Following a rectification process (Nnum 11, 2048 * 2048 to 935 * 924), we reconstructed light-field sequence into 3-D stacks (935 * 924 * 31 voxels) sequentially by LFD and VCD-Net and take the average reconstruction time of each frame as the criterion of throughput. As compared to the deconvolution method, the VCD-Net dramatically increases the reconstruction speed by over 900 times when the GPU was engaged, reaching a frame rate of 13.51 fps which is adequate for real-time reconstruction of dynamic processes.

| Computation Unit | Throughput of VCD-Net | Throughput of LFD |
|---|---|---|
| GPU (Nvidia GeForce RTX 2080Ti) | 0.074s / 13.51fps | 68.26s / 0.015 fps |
| CPU (Intel(R) Core(TM) i7-5930K) | 3.11s / 0.32 fps | 500.80s /0.0020fps |

**Supplementary Video 1.** 3D visualization of acting *C. elegans* (all neurons labelled) by VCD-LFM.
**Supplementary Video 2.** 3D visualization of acting *C. elegans* (A- and B- motor neurons labelled) by VCD-LFM.
**Supplementary Video 3.** Analysis of neural $Ca^{2+}$ signals and motion pattern in the acting *C. elegans.*
**Supplementary Video 4.** 3D visualizations of flowing red blood cells by LFD and VCD-LFM.
**Supplementary Video 5.** 3D visualizations of beating cardiomyocytes nuclei by LFD and VCD-LFM.
**Supplementary Video 6.** 3D visualizations of beating trabecular myocardium by LFD and VCD-LFM.
**Supplementary Video 7.** Dual-color 3D visualization revealing the dynamic process of red blood cells flowing through the tail vessels of the embryonic zebrafish.
**Supplementary Video 8.** 3D velocity distribution of the blood flow in the fish tail vessels.
**Supplementary Video 9.** Dual-color 3D visualization revealing the highly dynamic process of blood cells flowing in the beating zebrafish heart during cardiac cycles.
